# Global Mapping of circRNA-Target RNA Interactions Reveals P-Body-Mediated Translational Repression

**DOI:** 10.1101/2025.11.16.688748

**Authors:** Peng Li, Hongmei Zhang, Zhaokui Cai, Yuyang Zhang, Ruiyun Yang, Rong Ye, Jingxin Li, Hailian Zhao, Bowen Liu, Zhen Yuan, Xuekun Li, Xi Wang, Ping Zhu, Yuanchao Xue

**Author notes:** These authors contributed equally (Peng Li, Hongmei Zhang, Zhaokui Cai, Yuyang Zhang). **Corresponding author:** Yuanchao Xue.

## Abstract

Circular RNAs (circRNAs) are primarily produced through pre-mRNA back-splicing, yet their target repertoire and functional mechanisms remain elusive. Here, we present circTargetMap, a computational framework for globally mapping circRNA targets using RNA-RNA interactomes obtained via RNA in situ conformation sequencing (RIC-seq) in the hippocampus and ten human cell lines. This approach identified 117,163 high-confidence circRNA-target RNA interactions, with 83% of target mRNAs bound by multiple circRNAs. Functionally, *CDR1as* and *circRMST* repress target mRNA translation by sequestering them into processing bodies (P-bodies)—membraneless granules—through sequence-specific base-pairing, probably independent of AGO2, DICER, and miRNAs. To directly capture granule-associated interactions, we developed granule RIC-seq (GRIC-seq) method, revealing the broad role of circRNA-target RNA interactions in translational repression. Moreover, pathogenic variants are significantly enriched around circRNA-target RNA interaction sites, suggesting potential roles in disease. Our study provides valuable resources for circRNA functional exploration and a framework for investigating RNA-RNA interactions within membraneless organelles.

## INTRODUCTION

Circular RNAs (circRNAs) are covalently closed RNA molecules generated primarily through pre-mRNA back-splicing, which joins a downstream splice donor to an upstream acceptor site to form a characteristic back-splicing junction (BSJ)^1–4^. This process gives rise to thousands of distinct circRNAs across diverse genes^5–7^. Among these, a subset with high circular-to-linear expression ratios—such as *CDR1as*^8,9^(also known as *ciRS-7*), *circRMST*^10,11^, and *circHIPK3*^12^—are exceptionally stable, abundantly expressed, and consistently detected across cell types and tissues, suggesting evolutionarily conserved regulatory roles. CircRNAs also exhibit cell-type, tissue, and developmental stage-specific expression patterns^13,14^, with exceptionally high abundance in the brain^10,15,16^.

Although circRNAs have been implicated in key biological processes, including differentiation, cancer, and immune regulation^17–26^, their target repertoire and mechanisms of action remain poorly understood. Current models propose several functions for circRNAs, including acting as molecular sponges for microRNAs (miRNAs)^8,9^ and sequestering RNA-binding proteins (RBPs)^24^. The best-known example, *CDR1as*, contains over 70 conserved binding sites for *miR-7*, modulating its availability^8,9,27^. In addition, circRNAs also appear to be involved in the regulation of transcription and splicing^28,29^. However, these functions are primarily derived from isolated case studies, and it is unclear whether a broader, generalizable mechanism exists.

Given their single-stranded nature and stability, circRNAs may also function via direct base-pairing with target RNAs. Recent studies explored this possibility using 4′-aminomethyl-4,5′,8-trimethylpsoralen (AMT)-mediated psoralen crosslinking to capture circRNA-mRNA duplexes, followed by oligo pulldown and high-throughput RNA sequencing^30,31^. These approaches identified hundreds of mRNA-interacting circRNAs and several *circZNF609*-target RNA pairs but were limited by low resolution, lack of precise binding site information, and reliance on labor-intensive, pairwise validation. Bioinformatic predictions of circRNA-mRNA binding sites have also been attempted^30,31^, but a scalable, high-resolution approach for systematically mapping circRNA-target interactions is still lacking.

Here, we present circTargetMap, a computational framework that globally maps circRNA-target RNA interactions by analyzing RIC-seq data^32,33^—a technique that profiles the native RNA-RNA interactomes mediated by diverse RBPs. Applying this framework to data from ten cell lines and human/mouse hippocampus, we identified 117,163 high-confidence interactions. We show that *CDR1as* and *circRMST* repress translation of their targets via direct base-pairing—independently of AGO2, DICER, or miRNAs—by sequestering them into processing bodies (P-bodies), a class of membraneless granules^34^. To map these interactions within granules, we developed granule RIC-seq (GRIC-seq), enabling transcriptome-wide detection of circRNA-mRNA interactions in P-bodies. This revealed a widespread, P-body-mediated mechanism of circRNA-dependent translational repression. Furthermore, pathogenic variants are significantly enriched around circRNA-target junctions, implicating disrupted RNA-RNA interactions in disease. Together, our work uncovers a previously unrecognized mode of post-transcriptional regulation by circRNAs and establishes a generalizable framework for exploring RNA-RNA interactions in membraneless compartments, with broad implications for biotechnology and human health.

## RESULTS

### Global mapping of circRNA-target RNA interactions

CircRNAs are primarily generated by back-splicing events of pre-mRNAs^1–4^. Although numerous circRNAs have been identified, a transcriptome-wide map of circRNA-target RNA interactions is still lacking. To systematically identify the circRNA-target RNA interactions, we constructed RIC-seq libraries from human and mouse hippocampal tissues, human neural progenitor cell (hNPC)-derived neurons (using a well-established differentiation protocol^35,36^), human embryonic kidney cells (HEK293T), and the human colorectal adenocarcinoma cells HT29. Alongside our previously generated RIC-seq libraries from GM12878, HeLa, HepG2, IMR90, K562, H1-hESC, and hNPCs^32,37^, we obtained a total of 5.96 billion uniquely mappable reads, with an average chimeric read (reads that align to different positions within a single RNA or to different RNAs) ratio of 13.33% (Table S1). To estimate the reliability of RIC-seq captured RNA-RNA interactions, we also extracted RNA fragments before proximity ligation in neurons for subsequent *in vitro* random ligation and sequencing (Figure S1A). Only 1.7% of the chimeric reads identified in the *in situ* proximity ligation libraries overlapped with those from *in vitro* random ligation libraries (Figure S1B), indicating that RIC-seq efficiently captures genuine RNA-RNA interactions with minimal false positives.

Given that most circRNAs are expressed at lower levels than linear mRNAs and are sequence identical with their linear counterparts, except at the BSJ site, we developed a computational strategy, circTargetMap, to screen for chimeric reads that span the BSJ of circRNAs and simultaneously map to target RNAs (Figure 1A; see Methods). In theory, these chimeric reads represent potential circRNA-target RNA interactions. We subsequently employed a Monte Carlo simulation^38^ to identify high-confidence circRNA-target RNA interactions by comparing the chimeric read counts of observed pairwise interactions with those from simulated random interactions (*n* = 10^5^, *P* < 0.05; see Methods; Figure 1A). These efforts identified 5,840 high-confidence interactions across all analyzed cell lines and tissues, ranging from 24 in HEK293T cells to 1,435 in hNPC-derived neurons (Figure 1B; Table S2).

**Figure 1.**
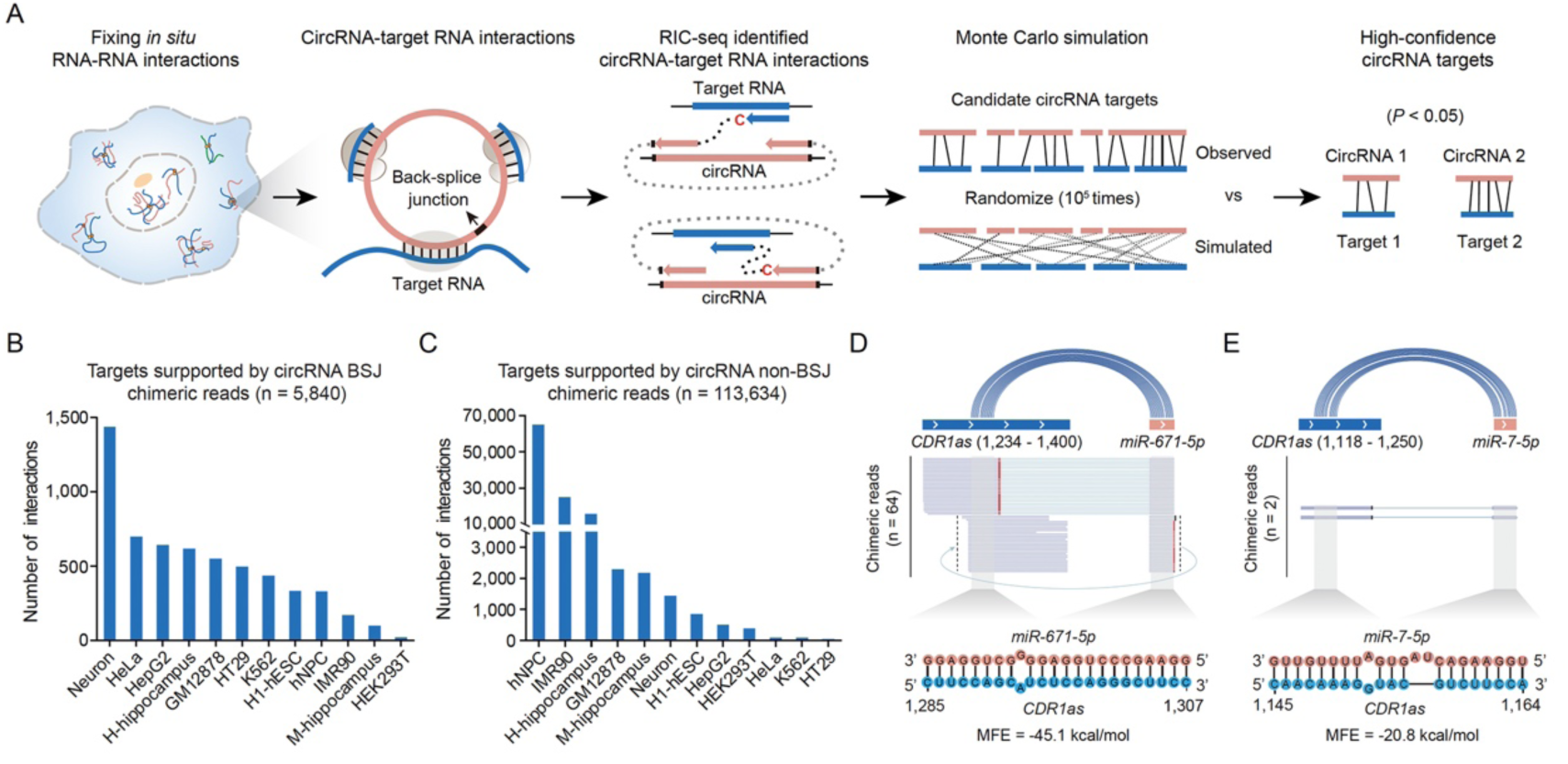
Global mapping of circRNA-target RNA interactions based on RIC-seq data. (A) Diagram of the RIC-seq technique and circTargetMap computational strategy for profiling spatial interactions between circRNAs and target RNAs. (B) Bar plot showing circRNA-target RNA interactions identified via BSJ sites across indicated cell lines and hippocampal tissues. (C) Bar plot showing circRNA-target RNA interactions identified using the entire circRNA sequence. CircRNAs with a circular-to-linear ratio ≥ 0.85 across the indicated cell lines and tissues were analyzed. (D-E) RIC-seq recapitulates known interactions: *CDR1as* with *miR-671-5p* (D) and *miR-7-5p* (E). Gray lines: chimeric reads; red/black lines: read junctures (red highlights pCp insertions). Lower panels: base-pairing regions between *CDR1as* and miRNAs.

While BSJ-containing chimeric reads effectively eliminate false-positive interactions from linear counterparts, this approach cannot efficiently identify targets of circRNAs with extremely high circularization ratios (≥0.85), such as *CDR1as* (*hsa_circ_0001946*). We therefore modified our strategy to incorporate all chimeric reads mapped across entire circRNAs for highly circularized species, rather than exclusively relying on BSJ-spanning reads. Using this revised approach (see Methods), we identified 113,634 high-confidence interactions across all examined cell lines and tissues, ranging from 5 in HT29 cells to 64,865 in hNPCs (Figure 1C; Table S3). Notably, our modified approach faithfully recapitulates well-known base-pairing interactions between *CDR1as*–*miR671-5p* and *CDR1as*–*miR7-5p* (Figures 1D and 1E).

By integrating BSJ and non-BSJ targets, we identified 117,163 high-confidence circRNA-target RNA interactions between 4,822 circRNAs and 16,474 target RNAs across all samples. Among these, 1,539 circRNAs interacted with more than five target RNAs, including 500 circRNAs that exhibited broad targeting capacity with over 50 RNA targets each (Figure S1C). Notably, we identified 7,176 target RNAs bound by more than five circRNAs each, including 115 RNAs targeted by over 50 circRNAs (Figure S1D). These results demonstrate that circRNAs form extensive and highly interconnected regulatory networks with target RNAs.

Analysis of these interactions revealed that target sites were preferentially located in coding sequences (CDS, 35.7%), followed by introns (27.8%), 5’UTR (15.6%), and 3’UTR (13.7%) (Figure S1E). To assess their functional relevance, we examined evolutionary conservation and found that circRNA-interacting regions on target RNAs were significantly more conserved than randomly shuffled regions (Figure S1F), suggesting these interactions may be biologically important. Furthermore, using the DuplexFold algorithm^39^, we calculated base-pairing potential and observed that interacting circRNA-target RNA fragments had lower minimum free energy (MFE) than shuffled controls (Figure S1G), supporting the involvement of RNA-RNA base pairing in mediating these interactions.

### Validation of circRNA-mRNA interactions

We next focused on circRNA-mRNA interactions, as circRNA mainly bound exonic regions of protein-coding genes (Figure S1E). To validate the identified circRNA-mRNA interactions, we co-stained circRNAs with their targets using hybridization chain reaction fluorescence *in situ* hybridization (HCR-FISH)^40^ in hNPC-derived neurons (Figures S2A and S2B). For spatial proximity analysis, we selected three representative circRNAs spanning a wide expression range: *CDR1as* (Counts in Per Million mapped reads (CPM) = 131.03 in human hippocampus; 96.24 in hNPC-derived neurons), *circRMST* (*hsa-RMST_0004*, 91.08 in neurons), and *circMAN1A2* (*hsa_circ_0000118*, 0.78 in neurons). From their interacting mRNAs, we randomly chose *MAN2A1* (5.19 in human hippocampus), *MTURN* (9.71 in human hippocampus), *FBH1* (2.69 in neurons), and *RARS2* (1.79 in neurons, BSJ targets for *CDR1as*); *APC2* (13.16 in neurons), *PALS2* (0.17 in neurons), and *CYB5R4* (0.2 in neurons, BSJ targets for *circRMST*); and *PTPRM* (0.98 in neurons) and *MAP3K2* (0.92 in neurons, BSJ targets for *circMAN1A2*) for HCR-FISH validation (Figures 2A and S2C). In addition, three non-BSJ supporting targets, *GRIN2B* (CPM = 0.84 in neurons), *MYO5A* (1.68 in neurons), and *AKAP9* (0.83 in neurons) for *CDR1as*, were selected for smFISH validation (Figure 2C). As a negative control, we used *POLR2C* mRNA, which does not interact with any of these three circRNAs.

**Figure 2.**
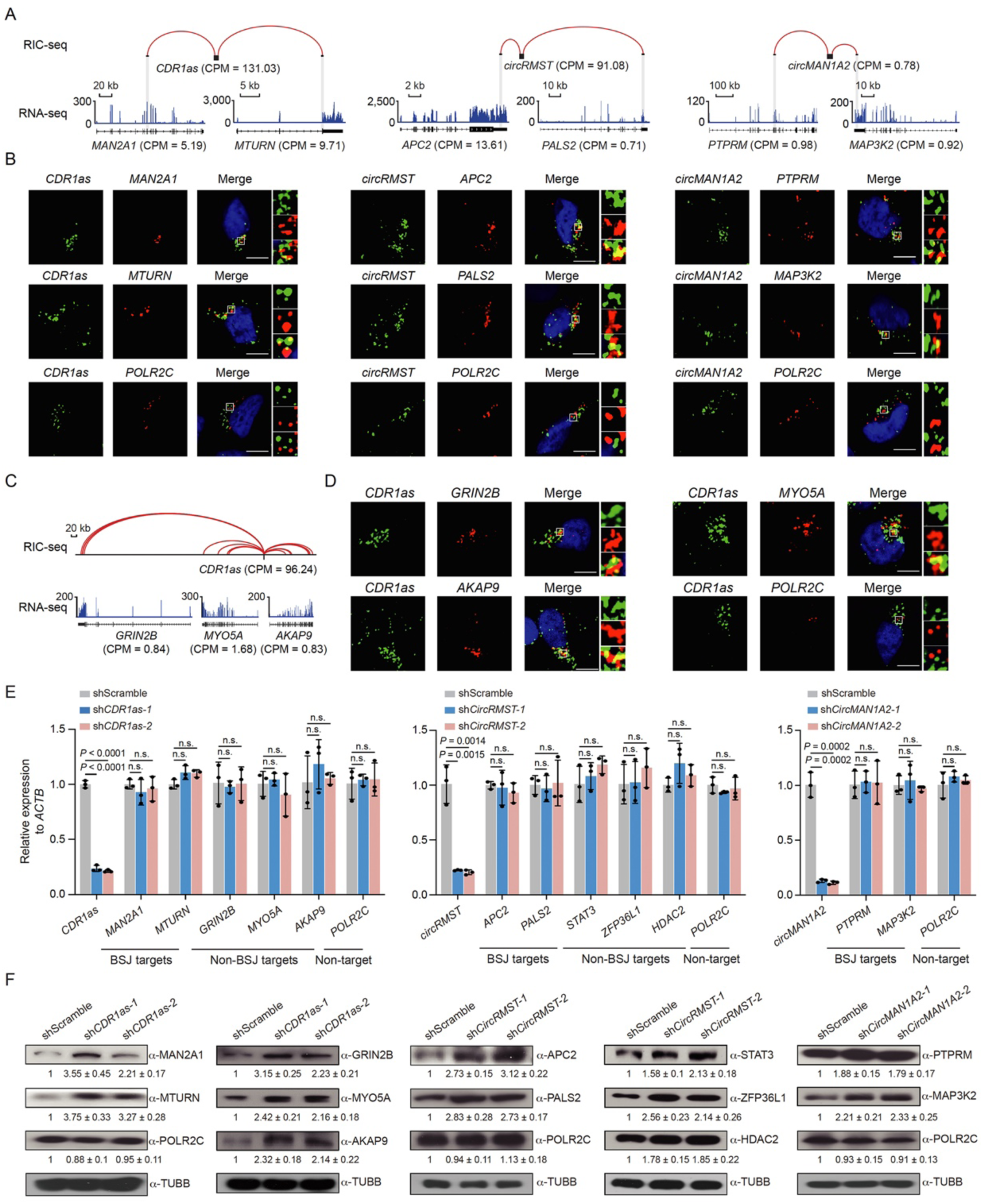
Validation of circRNAs-mRNA interactions. (A) RIC-seq and RNA-seq tracks showing interactions of *CDR1as* and its target mRNAs in the human hippocampus, and between *circRMST* or *circMAN1A2* with their targets in hNPC-derived neurons. Expression levels are presented as CPM. (B) HCR-FISH showing colocalization of *CDR1as*, *circRMST*, and *circMAN1A2* with their target mRNAs in neurons, but not with the non-target control *POLR2C*. DAPI (blue), circRNA (green), and mRNAs (red). Scale bar: 5 μm. (C) RIC-seq and RNA-seq tracks showing *CDR1as* interactions with *GRIN2B*, *MYO5A*, and *AKAP9* in neurons. (D) smFISH showing colocalization of *CDR1as* with target mRNAs in neurons, but not with *POLR2C*. Scale bar, 5 μm. (E) qPCR analysis of circRNAs and target mRNAs following circRNA knockdown in neurons. *POLR2C* served as a non-target control. BSJ targets: identified via the circRNA BSJ site. Non-BSJ targets: identified via the entire circRNA sequence. Data are presented as mean ± s.d.; *n* = 3 independent replicates, two-tailed unpaired Student’s *t*-test. *n.s.*, non-significant. (F) Immunoblotting analysis demonstrates increased protein expression of circRNA targets after circRNA knockdown in neurons. *POLR2C* served as a non-target control, and TUBB as a loading control. Band intensities were normalized to scramble shRNA (shScramble) treated samples (*n* = 3 independent replicates).

The FISH probes showed high target specificity, as evidenced by near-complete signal loss following knockdown of *CDR1as*, *circRMST*, and *circMAN1A2* (Figure S2D). HCR-FISH and smFISH analyses demonstrated that all these circRNAs colocalize with their target mRNAs in the cytoplasm of hNPC-derived neurons, while no colocalization was observed for the negative control *POLR2C* (Figures 2B, 2D, and S2E). Quantitative analysis showed that 55–74% of the target signals colocalized with their corresponding circRNAs in over 85% of neurons (*n* = 50, Figure S2F), supporting the specificity and robustness of the circRNA-target RNA interactions identified by our approach.

To validate the regulatory role of circRNAs on their targets, we knocked down the three selected circRNAs in hNPC-derived neurons using two different lentivirus-delivered short-hairpin RNAs (shRNA) targeting the BSJ site (see Methods). Following lentiviral infection, the expression levels of these three circRNAs were reduced by over 75% compared to scramble controls (Figure 2E), without affecting their linear RNA counterparts (Figure S2G). Surprisingly, although circRNA depletion did not alter the mRNA levels of any examined BSJ or non-BSJ targets (Figure 2E), it consistently increased the protein levels of all 12 tested targets by at least 1.6-fold (Figure 2F). These findings suggest that these circRNAs repress the translation of their target mRNAs.

### *CDR1as* globally represses target mRNA translation

To evaluate whether circRNA knockdown globally enhances target mRNA translation, we selected *CDR1as* for transcriptome-wide analysis due to its high abundance and numerous targets. Paired Smart-seq2 ^41^ (low-input RNA-seq) and Ribo-lite^42^ (low-input Ribo-seq) analyses of *CDR1as*-depleted neurons showed high reproducibility between replicates (*R* > 0.97, Figures 3A, S3A, and S3B). Among 1,036 neuronal targets of *CDR1as*, 863 were detected by both RNA-seq and Ribo-seq. Of these, only 19 showed significant RNA-level changes after knockdown (|fold change| > 1.5, *P* < 0.05; Figure 3B; Table S4), suggesting that *CDR1as* has a minimal impact on mRNA abundance.

**Figure 3.**
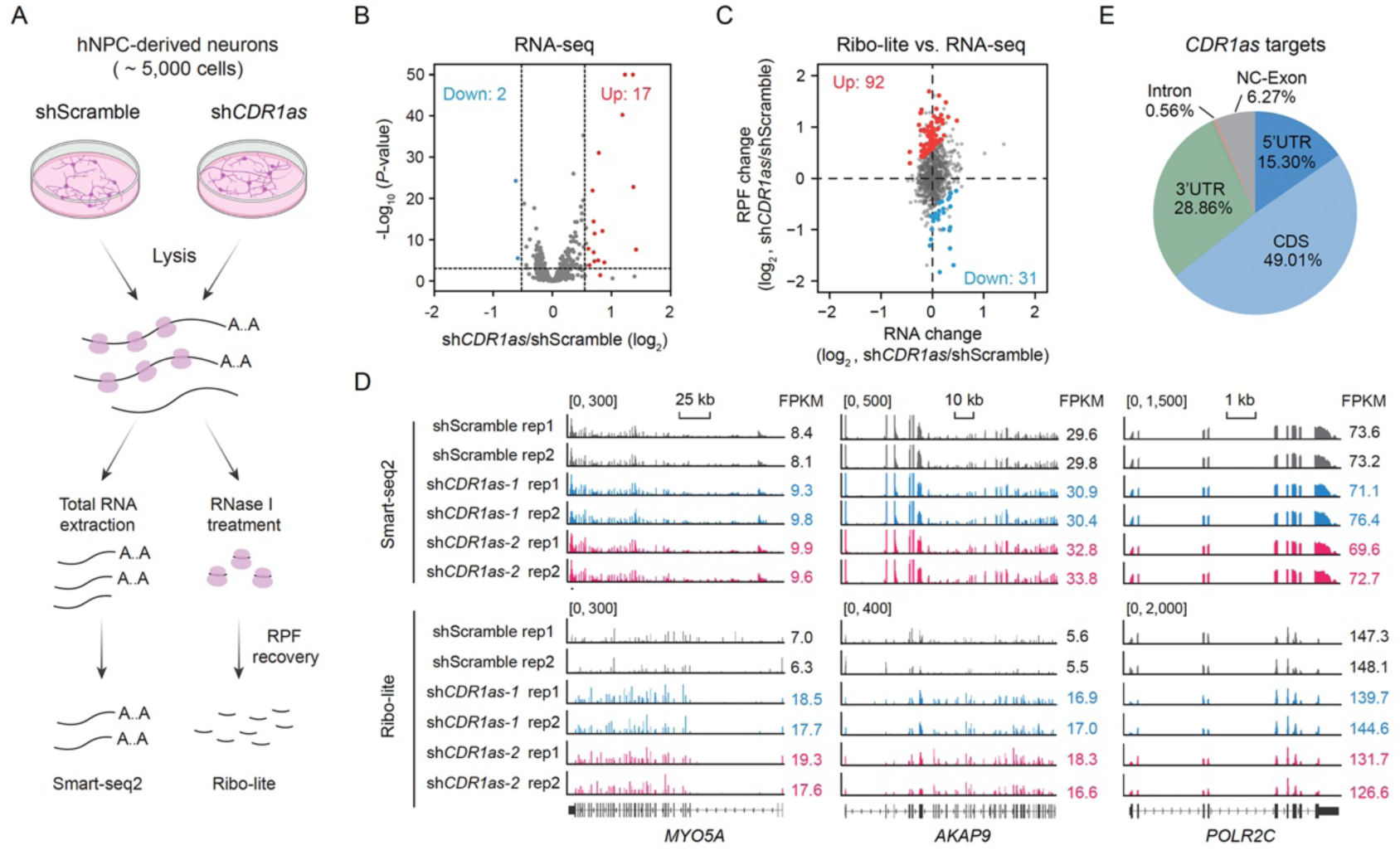
*CDR1as* globally represses target mRNA translation. (A) Schematic of Smart-seq2 and Ribo-lite analyses to detect mRNA and RPF changes upon *CDR1as* knockdown in 5,000 neurons. (B) RNA level changes of *CDR1as* targets upon *CDR1as* knockdown in neurons. Red: upregulated, blue: downregulated. (C) Scatter plots comparing mRNA (x-axis) and RPF (y-axis) changes of *CDR1as* targets with unchanged mRNA levels. (D) Smart-seq2 (mRNA) and Ribo-lite (RPF) tracks of target mRNAs (*MYO5A*, *AKAP9*) and non-target *POLR2C* after *CDR1as* knockdown. Rep, replicates. (E) Pie chart showing the distribution of *CDR1as* interaction sites across target RNA regions, normalized to total genomic length. NC-Exon, exon of noncoding RNA.

To determine whether *CDR1as* regulates translation efficiency (TE) – defined as the ratio of ribosome-protected fragments (RPFs) to mRNA abundance – we integrated Ribo-seq and RNA-seq data. Intriguingly, while mRNA levels of 844 *CDR1as* targets remained unchanged upon *CDR1as* knockdown, TE was significantly altered for 123 of them, with 92 increased and 31 decreased (|FC| > 1.5, p < 0.05; Figure 3C; Table S4). Notably, *CDR1as* expression was remarkably high, exceeding target RNA abundance by 15.1-fold on average (Figure S3C). Binding intensity analysis revealed stronger interactions of *CDR1as* with up-regulated targets compared to unchanged targets (Figure S3D). Consistent with the results from qPCR and immunoblotting (Figures 2E and 2F), two *CDR1as* targets, *MYO5A* and *AKAP9*, exhibited no changes in RNA levels according to RNA-seq, yet their RPF signals increased by at least 2.7-fold based on Ribo-seq data (Figure 3D). As a non-target control, *POLR2C* showed negligible changes in both mRNA and RPF signals. This translational repression function aligns with *CDR1as*’s preferential binding to CDS and 3’UTR regions (Figure 3E), which are known to play critical roles in translation regulation. These findings demonstrate that *CDR1as* globally represses the translation of its interacting mRNAs in hNPC-derived neurons.

### AGO2/miRNA-independent translational repression by *CDR1as*

While *CDR1as* has been reported to regulate gene expression by sponging *miR-7* in brain tissue^27^, we investigated whether its translational repression requires AGO2 or miRNAs. Using *AGO2* knockout (KO, miRNA-function impaired)^43^ and *DICER* KO (mature miRNA-deficient)^44^ 293T cells, we assessed *CDR1as*-mediated translational repression using psiCHECK-2 dual-luciferase reporters containing target sequences (*GRIN2B*/*MYO5A* 3’UTRs, or *AKAP9* CDS; Figures S4A and S4B). Upon individually transfecting these luciferase reporter plasmids into *CDR1as* overexpressing (80% circularization efficiency) *AGO2*-KO, *DICER*-KO, or wild-type (WT)-293T cells (Figures S4C and S4D), the relative luciferase activity (Renilla/Firefly) of all three targets decreased in a *CDR1as* dose-dependent manner across all three genetic backgrounds (Figure S4E). In contrast, no changes were observed for the negative control luciferase reporter containing an equal length of lambda DNA (γDNA) fragment (Figure S4E). Importantly, all the reporter RNA levels were unchanged (Figure S4F). These results demonstrate that *CDR1as* likely represses target mRNA translation independent of AGO2 and miRNAs.

### *CDR1as* and *circRMST* repress target mRNA translation through base pairings

Next, we examined whether *CDR1as* represses the translation of its target mRNAs through base pairing. To this end, we extracted the chimeric interacting RNA fragments and deduced RNA duplexes using the RNA DuplexFold algorithm^39^ for *CDR1as* and each interacting target mRNA fragment (*GRIN2B, MYO5A*, and *AKAP9*) derived from the non-BSJ regions (Figures 4A, 4D, and 4G). Using dual-luciferase reporters containing these fragments (Figure S4B), we systematically mutated specific binding segments (denoted as P1, P2, or P3) in these mRNAs to disrupt their base-pair potential with *CDR1as* (see the mutated sequences above each duplex), which was confirmed by reduced MFEs in the duplexes (Figures S5A-C). To further establish causality, we designed compensatory mutations in *CDR1as* that restored duplex formation with the corresponding mutant target mRNAs (Figures S5A-C).

**Figure 4.**
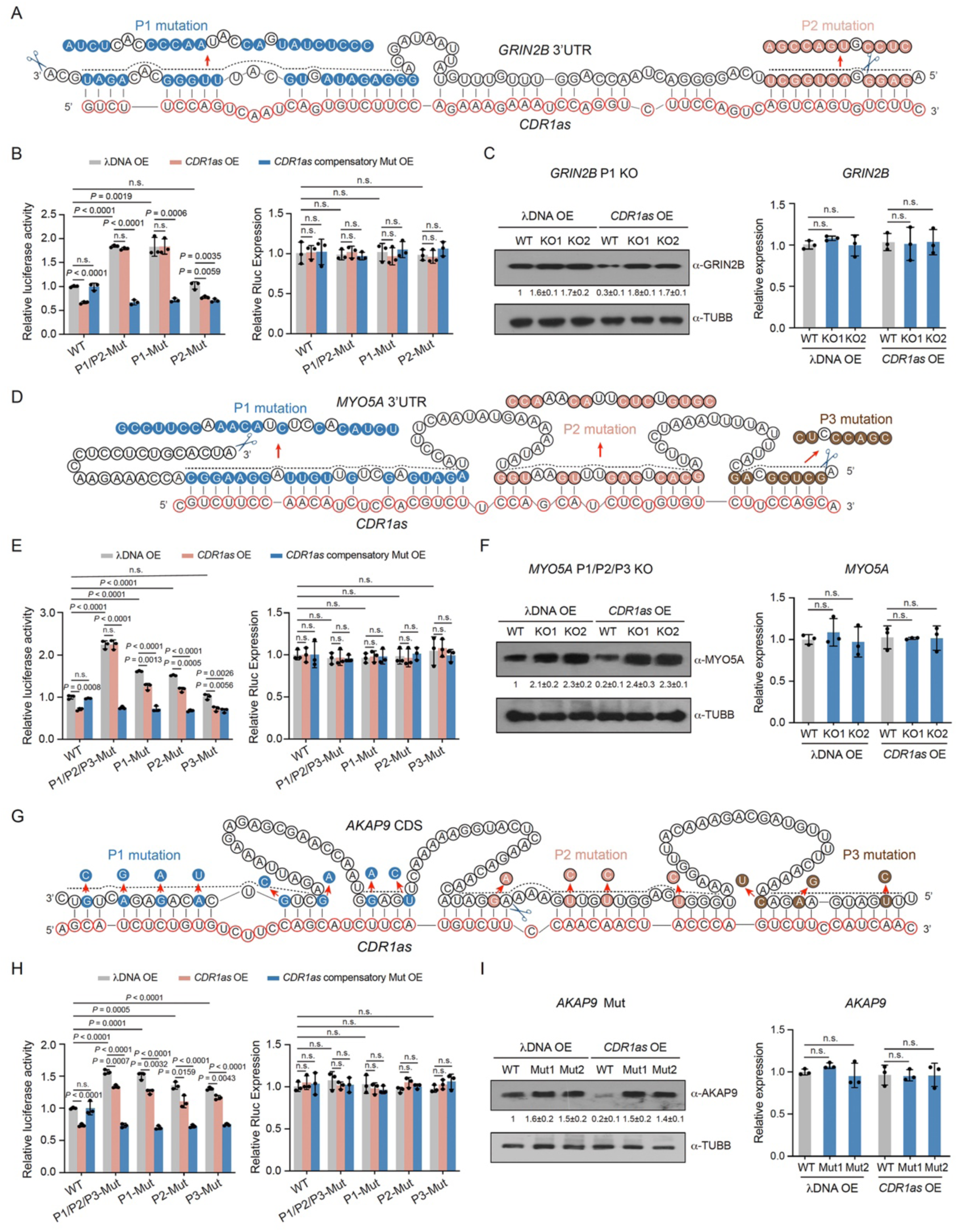
*CDR1as* represses target mRNA translation via base pairings. (A) Predicted duplex between *CDR1as* (internal region) and *GRIN2B* 3′UTR; arrows mark mutation sites. P1 (blue), P2 (pink); scissors indicate CRISPR-Cas9 cleavage site. (B) Luciferase assays showing relative activities (left) and mRNA levels (right) of *GRIN2B* 3 ′ UTR reporters (WT, P1/P2-Mut, P1-Mut, P2-Mut) in 293T cells overexpressing *CDR1as* or compensatory mutants. γDNA, overexpression control. (C) Immunoblotting (left) and qPCR (right) showing *GRIN2B* protein and RNA levels in WT and P1 KO 293T cells after *CDR1as* overexpression. (D–F) Similar analyses for *MYO5A* 3′UTR: duplex structure (D); luciferase reporter assays for WT and P1/P2/P3-Mut reporters (E); and protein/RNA levels in WT and KO cells (F). (G–I) Similar analyses for *AKAP9* CDS: duplex structure (G); luciferase assays for WT and synonymous mutant reporters (H); and protein/RNA levels in WT and mutant knock-in cells (I). Data in B, C, E, F, H, and I represent mean ± s.d. (*n* = 3 independent replicates); two-tailed unpaired Student’s *t*-test; *n.s.*, non-significant. Band intensities were normalized to λDNA-overexpressing WT cells.

For *GRIN2B* 3’UTR, mutation of the P1 segment eliminated the translational repression mediated by *CDR1as*—an effect observed in both WT and P2-Mut reporters—without affecting mRNA levels in *CDR1as*-overexpressing 293T cells (Figure 4B). Importantly, this loss of repression was rescued by introducing a corresponding compensatory mutation in *CDR1as* that restored base-pairing (Figure 4B). Due to technical challenges in achieving CRISPR-Cas9-mediated deletion of the P1 segment in hNPCs—attributed to low transfection efficiency—we used 293T cells as a surrogate model to investigate its endogenous function. This cell line exhibits transcriptomic similarity to neurons^45^ and expresses high endogenous levels of *CDR1as*^46,47^. Genetic deletion of the P1 segment in the *GRIN2B* endogenous locus resulted in a >50% increase in protein expression without altering mRNA levels (right panel, Figure 4C).

Similarly, analysis of *MYO5A* 3’UTR mutations (P1/P2/P3) showed that P1 and P2 segments were required for *CDR1as*-mediated translational repression, and this repression was rescued by corresponding compensatory mutations in *CDR1as* (Figure 4E). Furthermore, combinatorial deletion of P1 and P2 segments in the endogenous *MYO5A* locus led to a >100% increase in protein expression (left panel, Figure 4F). For *AKAP9*, synonymous CDS mutations that disrupt *CDR1as* binding elevated baseline translation, although residual repression indicates the presence of additional, weaker interactions. Compensatory mutations in *CDR1as* restored repression (Figure 4H), and knock-in of these mutations increased endogenous AKAP9 protein level by 50% and abolished *CDR1as*-mediated repression (Figure 4I). Together, these results demonstrate that *CDR1as* requires direct base-pairing with target mRNAs (*GRIN2B* via P1; *MYO5A* via P1/P2; *AKAP*9 via multiple CDS sites) to mediate translational repression without affecting mRNA abundance.

We next asked whether the circular form of *CDR1as* is required for this repression. Ectopic expression of circular, linear, or circularization-deficient *CDR1as* in HeLa cells (Figure S5D), which lack endogenous *CDR1as*^8^ expression, revealed that only the circular *CDR1as* significantly suppressed translation of Rellina luciferase reporters containing *GRIN2B 3’UTR*, *MYO5A 3’UTR*, or *AKAP9 CDS*, without changing their mRNA levels (Figures S5E-G). This repression was abolished when cognate binding sites in target RNAs were mutated (Figures S5E-G). Rescue experiments in *CDR1as*-knockdown 293T cells further confirmed that only circular *CDR1as* effectively restored the translational repression, while linear and circularization-deficient forms did not (Figures S5H-K). These findings suggest that the translational repression is strictly dependent on the circular topology of *CDR1as*.

This regulatory mechanism also applies to other circular RNAs, as demonstrated by *circRMST*. Mutations in the *circRMST* binding sites within the 3’UTRs of *APC2* and *PALS2* (BSJ-supported targets, Figures S6A and S6B) abolished *circRMST*-mediated repression in luciferase assays without altering mRNA levels in 293T cells, whereas compensatory mutations in *circRMST* effectively restored this repression (Figures S6C and S6D). Furthermore, only circular *circRMST*, not linear or circularization-deficient forms, effectively suppressed translation of reporters containing the *APC2* or *PALS2* 3’UTRs. This repression was abolished when the cognate binding sites in the target RNAs were mutated (Figures S6E-G). In summary, both *CDR1as* and *circRMST* suppress target mRNA translation through direct base-pairing without altering mRNA abundance, and their circular topology is essential for this translational repression.

### *CDR1as* and *circRMST* repress translation by sequestering target mRNAs into P-bodies

To elucidate the mechanism of *CDR1as*-mediated translational repression, we performed ChIRP-MS^48^ in hNPC-derived neurons to identify *CDR1as*-associated proteins (Figure S7A). Using *CDR1as*-specific probes, we achieved selective enrichment of endogenous *CDR1as* but not the control *ACTB* mRNA, compared to LacZ probes (Figure S7B). ChIRP-MS analysis identified 104 *CDR1as*-associated proteins, including 29 RBPs and 8 known P-body components^49,50^ (*P* = 3.06e-6; Figures 5A and S7C; Table S5), suggesting P-body involvement in *CDR1as*-mediated translational repression. Immunoblotting validation confirmed specific enrichment of three candidate proteins: IGF2BP1, HNRNPU, and the canonical P-body protein DDX6 (Figure S7D). These findings suggest that *CDR1as-*mediated translational repression is likely through sequestering target mRNAs into P-bodies.

**Figure 5.**
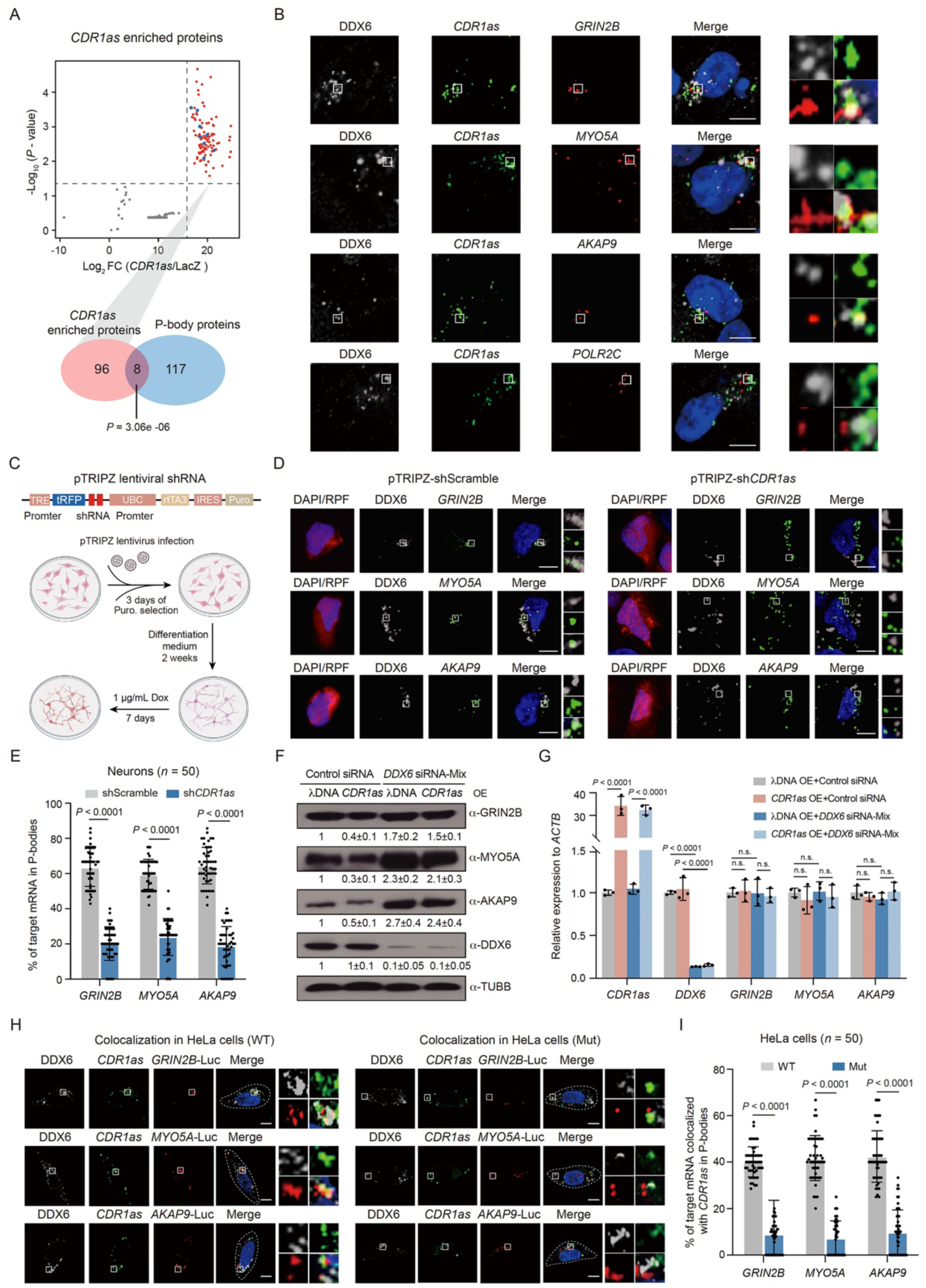
*CDR1as* sequesters target mRNAs into P-bodies for translational repression. (A) Scatter plot of *CDR1as*-interacting proteins identified by ChIRP-MS in neurons. Blue points, known P-body proteins. Venn diagram shows overlap between *CDR1as*-enriched and P-body proteins (one-tailed hypergeometric test). (B) IF/smFISH showing colocalization of *CDR1as* with target mRNAs (*GRIN2B, MYO5A,* and *AKAP9*) and P-body marker DDX6 in neurons, but not with non-target control *POLR2C*. DDX6, gray; DAPI, blue; *CDR1as*, green; mRNAs, red. Scale bar, 5 μm. (C) Schematic of the inducible shRNA knockdown strategy in neurons. (D, E) IF/smFISH and quantification showing reduced localization of target mRNAs in P-bodies upon *CDR1as* knockdown (tRFP+, red). Scale bar, 5 μm. (F, G) Immunoblotting (F) and qPCR (G) showing *CDR1as* targets and DDX6 expression after *DDX6* knockdown in *CDR1as*-overexpressing neurons. Data represent mean ± s.d. (*n* = 3); two-tailed unpaired Student’s t-test. Band intensities were normalized to control siRNA-treated neurons overexpressing the λDNA vector. (H, I) In *CDR1as*-overexpressing HeLa cells, *CDR1as* (green) colocalizes with DDX6 (gray) and Renilla mRNAs carrying WT, but not mutant, target fragments (red). Cell boundaries are outlined with white dashed lines; DAPI, blue. Scale bar, 5 μm. Quantification in (E) and (I) is presented as mean ± s.d.; *n* = 50 cells, two-tailed unpaired Student’s *t*-test.

To test this hypothesis, we performed IF/smFISH co-localization analysis in hNPC-derived neurons. *CDR1as* foci showed significant colocalization with both the P-body marker DDX6 and target mRNAs (*GRIN2B*, *MYO5A*, and *AKAP9*), but not the control mRNA *POLR2C* (Figures 5B and S7E). Using a doxycycline-inducible pTRIPZ shRNA system (Figure 5C), we achieved 80% knockdown of *CDR1as* in RFP-labeled neurons (Figure S7F). Subsequent IF/smFISH revealed markedly reduced target mRNA-DDX6 co-localization upon *CDR1as* depletion (Figures 5D and 5E), supporting a model where *CDR1as* sequesters target mRNAs into P-bodies for translational repression. Furthermore, knockdown of the core P-body components *DDX6* or *LSM14A* abolished *CDR1as* overexpression-mediated translational repression of *GRIN2B*, *MYO5A*, and *AKAP9* in hNPC-derived neurons (Figures 5F, 5G, S7G, and S7H), confirming that P-body integrity is required. Notably, IF/smFISH analysis of *DDX6*-depleted neurons showed that DDX6 loss did not affect the colocalization of *CDR1as* with its target mRNAs (Figure S7I).

To determine whether *CDR1as* requires base-pairing to localize target mRNAs into P-bodies, we transfected luciferase reporters containing either WT or mutated base-pairing sites (*GRIN2B*, *MYO5A*, and *AKAP9*) into *CDR1as-*ectopically expressed HeLa cells. Disruption of base-pairing significantly decreased the colocalizations between *CDR1as* and target mRNAs in P-bodies (Figures 5H and 5I), indicating that direct RNA-RNA base pairing is required for sequestration.

To explore RBPs involvement, we focused on IGF2BP1—one of eight *CDR1as*-interacting P-body proteins with characterized RNA-binding motifs^51^ and available PAR-CLIP data^52^. Motif analysis using FIMO^53^ revealed multiple significant motif occurrences within the *CDR1as* regions interacting with *GRIN2B*, *MYO5A*, and *AKAP9* (Figure S7J). Furthermore, public PAR-CLIP data^52^ confirmed IGF2BP1 binding at the *MYO5A* and *AKAP9* interaction sites in 293T cells (Figure S7J). Functionally, *IGF2BP1* knockdown in 293T cells markedly increased Rellina luciferase activity for all three targets, whereas re-expression of IGF2BP1 further restored the repression (Figures S7K-M). These findings demonstrate that IGF2BP1 may contribute to the *CDR1as*-mediated translational repression of target mRNAs.

Finally, we examined whether *circRMST* employs a similar mechanism. IF/HCR-FISH analysis in hNPC-derived neurons, revealing significant colocalization of *circRMST* foci with both DDX6 and target mRNAs (*APC2* and *PALS2*), but not the control mRNA *POLR2C* (Figures S8A and S8B). *cricRMST* knockdown in RFP-labeled neurons (80% efficiency, Figure S8C) led to a marked reduction in co-localization between target mRNAs and DDX6 (Figures S8D and S8E). Moreover, *DDX6* knockdown abolished the translational repression of *APC2* and *PALS2* mediated by *circRMST* overexpression, without affecting their RNA levels (Figures S8F and S8G). These results demonstrate that *circRMST* also represses translation by sequestering target mRNA into P-bodies via base-pairings, mirroring the mechanism employed by *CDR1as*.

### Systematic characterization of circRNA-mediated translational repression in P-bodies

To investigate whether circRNA-mediated translational repression is generally present in P-bodies, we reanalyzed RNA-seq data from fluorescence-activated particle sorting (FAPS) of GFP-LSM14A-marked P-bodies in HeLa cells^54^ (Figure 6A). This analysis identified 3,835 P-body-localized circRNAs, top-enriched including *hsa_circ_0000745* (*circSPECC1*), *hsa_circ_0128780* (*circGUSBP1*), and *hsa_circ_0001944* (Figure 6B; Table S6). Furthermore, among 686 circRNA targets identified in HeLa cells, 621 were mRNAs, of which 546 were detected by RNA-seq and showed significant enrichment in P-bodies compared to non-targets (Figure 6C).

**Figure 6.**
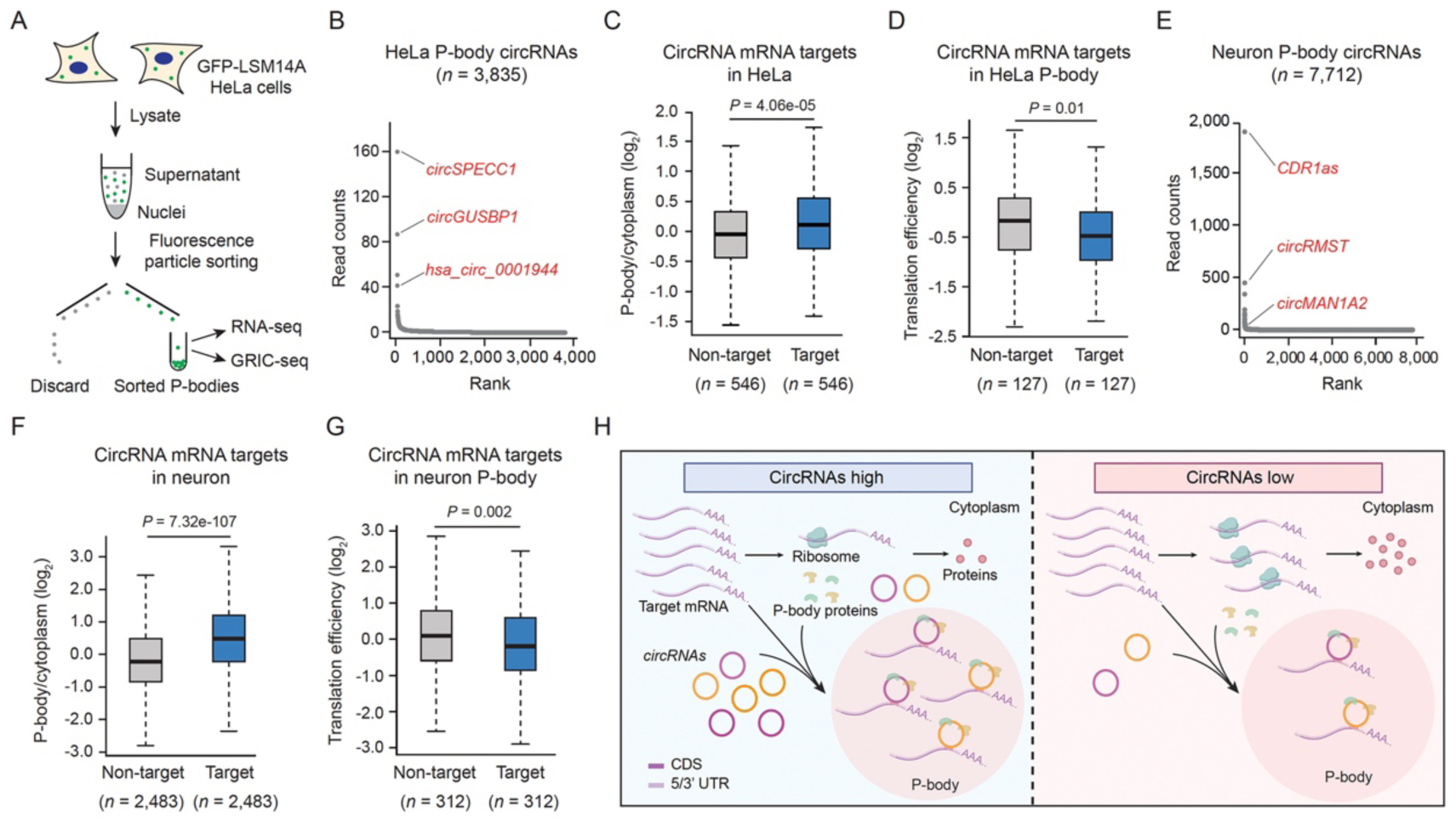
Widespread circRNA-mRNA interactions in P-bodies. (A) P-body purification via FAPS in GFP-LSM14A HeLa cells for subsequent GRIC-seq analysis. (B) Ranked abundance of detected circRNAs by RNA-seq in purified P-bodies from HeLa cells. (C) Boxplots showing the relative enrichment of circRNA target mRNAs compared with non-targets in HeLa cell P-bodies. (D) Boxplots showing the reduced TE of circRNA target mRNAs identified by GRIC-seq compared with non-targets in HeLa cells. (E) Ranked abundance of detected circRNAs by RNA-seq in purified P-bodies from hNPC-derived neurons. (F) Boxplots showing the relative enrichment of circRNA target mRNAs compared with non-targets in neuronal P-bodies. (G) Boxplot showing the reduced TE of circRNA target mRNAs identified by GRIC-seq compared with non-targets in neurons. The *P*-values in C, D, F, and G are calculated using a Wilcoxon rank-sum test. (H) Model illustrating circRNAs direct target mRNAs into P-bodies through sequence-specific base pairing, resulting in translation inhibition.

To directly identify circRNA-target RNA interactions within P-bodies, we developed a granule RIC-seq (GRIC-seq) method for global mapping of RNA-RNA interactions in FAPS sorted, GFP-LSM14A-labeled P-bodies from HeLa cells stably expressing GFP-LSM14A (see Methods). After separating nuclear and cytoplasmic fractions, we successfully isolated GFP-LSM14A–labeled P-bodies by comparing with GFP-LSM14A-Δ negative control (Figures S9A and S9B), as previously described^49,54^. This analysis revealed 147 high-confidence circRNA-target RNA interactions within these translational silencing compartments. Importantly, the TE of 127 target mRNAs was significantly lower than that of non-targets (Figure 6D), suggesting translational repression of these target mRNAs within P-bodies.

Next, we adopted an immunoprecipitation-based approach^55,56^ to identify P-body-enriched circRNAs in hNPC-derived neurons (Figure S9C). P-bodies were isolated from the cytoplasmic fraction of neurons using a specific antibody against LSM14A, followed by RNA-seq and GRIC-seq. Such analysis identified 7,712 P-bodies-localized circRNAs (Figure 6E; Table S7), including the examined *CDR1as*, *circRMST*, and *circMAN1A2*. Again, among the 3,010 neuronal targets of circRNAs, 2,668 were mRNAs, of which 2,483 were detected by RNA-seq and significantly enriched in P-bodies compared to non-targets (Figure 6F). This included validated targets (*GRIN2B*, *AKAP9*, and *MYO5A* for *CDR1as*; *STAT3* and *ZFP36L1* for *circRMST*) but not the non-target mRNA *POLR2C* (Figure S9D). GRIC-seq analysis in the LSM14A-immunoprecipitated P-bodies identified 384 circRNA-target RNA interactions, 333 were mRNAs, of which 312 were also detected by RNA-seq and Ribo-seq, showing significantly lower TE than non-targets (Figure 6G). These findings suggest a model wherein circRNAs base-pair with target mRNAs to direct their P-body sequestration and translational repression (Figure 6H).

### Disease-associated variants in circRNA-target RNA interaction sites

To investigate the pathological relevance of circRNA-target RNA interactions, we collected 78,298 disease-associated variants from the GWAS Catalog^57^ and 251,116 pathogenic or likely pathogenic variants from the ClinVar database^58^. Mapping these variants to circRNA-target RNA interaction regions revealed 14,548 overlapping variants, including 8,339 single-nucleotide variants (SNV), 4,557 deletions, 551 duplications, 550 microsatellite variants, 320 indels, and 231 insertions (Figure 7A). Notably, disease-associated variants showed significant enrichment around chimeric junctions compared to randomly shuffled controls (Figure 7B, see methods), suggesting potential roles of circRNA-target RNA interactions in disease pathogenesis. To identify functionally disruptive variants, we assessed their impact on RNA duplex stability by calculating the mutation-induced changes in minimum free energy (ΔMFE). Variants were classified as high-impact if they met both criteria: |ΔMFE| ≥ 3 kcal/mol and a ≥ 20% change in stability relative to the wild-type duplex. Using this threshold, we identified 27.12% (1,457 of 5,372) of circRNA-located variants and 15.24% (1,492 of 9,790) of target RNA-located variants as high-impact (Figure 7C).

**Figure 7.**
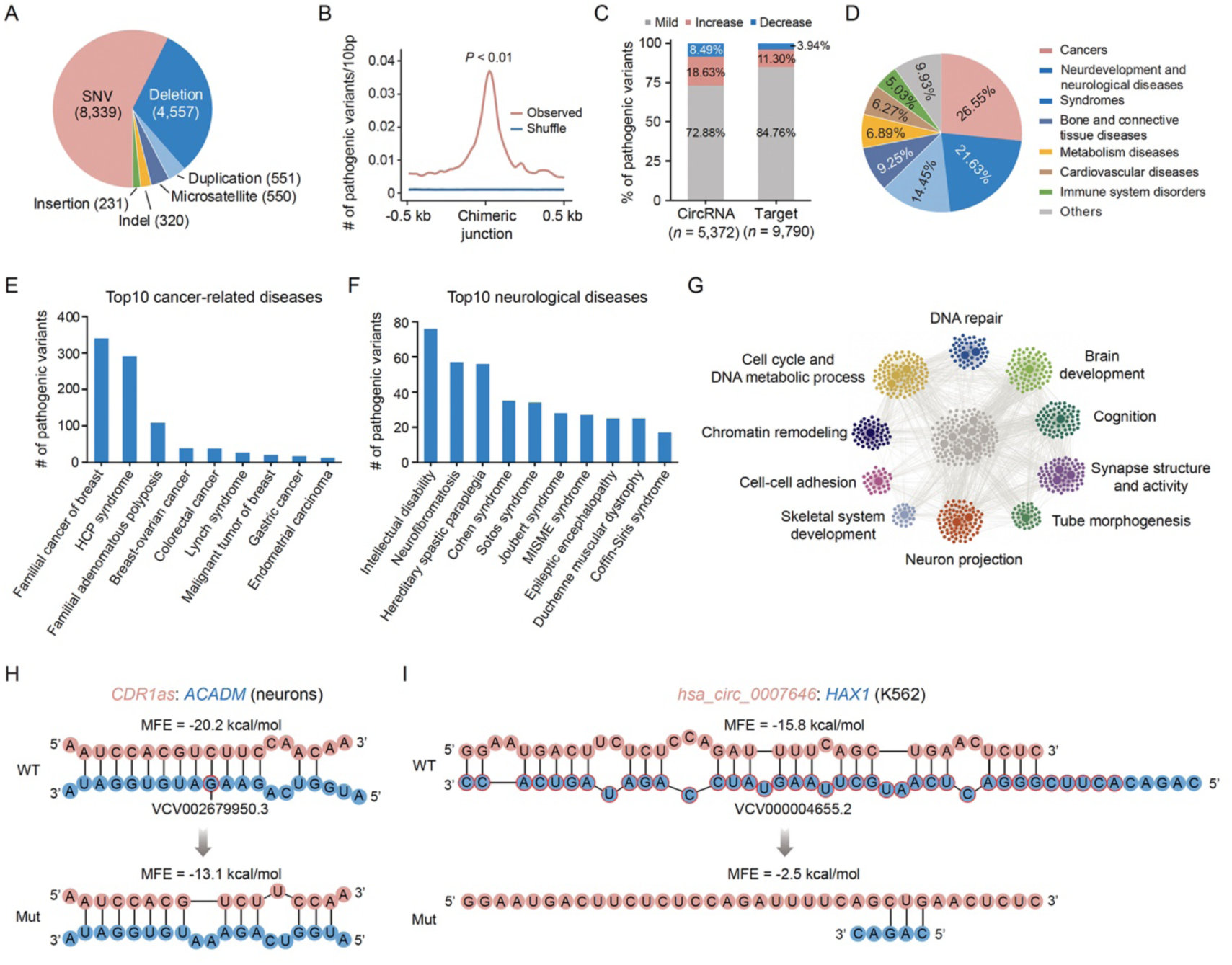
Disease-associated variants are enriched in circRNA-target RNA interaction regions. (A) Distribution of disease-associated variant types within circRNA-target RNA interaction sites. (B) Pathogenic variants are enriched around circRNA-target RNA interaction sites (pink) compared to randomly shuffled controls (blue). *p*-value from 100 randomizations. (C) Bar plot illustrating the proportion of variants affecting circRNA-target RNA interaction stability. “Increase” and “Decrease” denote |ΔMFE| ≥ 3 kcal/mol with ≥ 20% MFE increase or decrease, respectively; “Mild” denotes |ΔMFE| < 3 kcal/mol and <20% MFE change. (D) Pie chart of diseases linked to high-impact variants. (E, F) Number of high-impact variants in top 10 cancer-related (E) and neurological (F) diseases; HCP, hereditary cancer-predisposing; MISME, microcephaly-intellectual disability-sensorineural hearing loss-epilepsy-abnormal muscle tone. (G) GO enrichment of genes affected by high-impact variants. (H, I) RNA duplexes showing *CDR1as*–*ACADM* (H) and *hsa_circ_0007646*–*HAX1* (I) interactions in WT and mutant RNAs; red circles indicate nucleotide deletions.

Linking these high-impact variants to disease annotations revealed strong associations with cancers (26.55%), neurological disorders (21.63%), and genetic syndromes (14.45%) (Figure 7D). Among cancer subtypes, familial breast cancer exhibited the highest variant burden (340 variants), followed by hereditary cancer-predisposing (HCP) syndrome (291 variants) and familial adenomatous polyposis (109 variants) (Figure 7E). In neurological disorders, intellectual disability carried the largest number of variants (76 variants) (Figure 7F). Gene Ontology analysis of the affected target genes revealed significant enrichment in cancer-related pathways (*e.g.,* cell proliferation and DNA repair) and neuronal processes (*e.g.,* neuron projection and brain development) (Figure 7G). To mechanistically connect these variants to circRNA function, we highlight two examples where mutations disrupt base-pairing interactions. A pathogenic G deletion in *ACADM*, a neuronal target of *CDR1as*, weakens their binding (MFE from –20.2 to –13.1 kcal/mol) (Figure 7H), while a large deletion at the *hsa_circ_0007646* binding site in *HAX1* abolishes their interaction in K562 cells (Figure 7I). Together, these results suggest that disease-associated variants may contribute to pathogenesis by disrupting circRNA–mRNA base-pairing interactions.

## DISCUSSION

Although numerous circRNAs have been identified, systematically identifying their target repertoires still represents a grand challenge^59^. In this study, we developed a computational strategy, circTargetMap, to globally map circRNA-target RNA interactions using RIC-seq-identified *in situ* RNA-RNA interactomes from tissues and various cell lines. Our efforts led to the identification of the largest collection of circRNA-target RNA interactions. By characterizing these interactions, we unexpectedly found that *CDR1as*, *circRMST*, and *circMAN1A2* mainly function to repress translation rather than modulate RNA abundance. This translational repression mechanism is achieved by base pairing with target mRNAs and sequestering them into P-bodies. RNA-seq and GRIC-seq analysis of biochemically purified P-bodies further identified thousands of P-body-localized circRNAs and hundreds of circRNA-mRNA interactions within the translational silencing compartments.

Our study elucidates a potentially widespread mechanism of gene regulation whereby circRNAs directly repress target mRNA translation through sequence-specific base-pairing and P-body recruitment. Several key findings support this model: (1) *CDR1as* and other circRNAs physically interact with target mRNAs via defined base-pairing regions; (2) these interactions mediate translocation of target mRNAs into P-bodies; (3) the process occurs independently of canonical miRNA/AGO2 pathways; (4) approximately 83% of target mRNAs are bounded by multiple circRNAs; and (5) only circular isoforms possess this regulatory capacity. These results significantly expand our understanding of circRNA function beyond miRNA sponging, revealing a widespread translational regulatory network. This translational repression mechanism may prevent undesired mRNA translation under specific physiological conditions, for example, neurogenesis and synaptic signaling transmission processes in the hippocampus and neurons.

P-bodies are cytoplasmic, membraneless ribonucleoprotein (RNP) granules^60,61^. While initially characterized as sites of mRNA decay, recent evidence suggests P-bodies primarily serve as hubs for mRNA storage and translational repression^49,62^. In this study, we observed significant enrichment of circRNAs and their target mRNAs in P-bodies, accompanied by markedly reduced TE of these targets in both HeLa cells and neurons. These findings reveal P-body dependence as a crucial mechanistic feature of circRNA-mediated gene regulation. Through ChIRP-MS and immunoblotting, we identified specific interactions between *CDR1as* and core P-body components. Imaging studies further demonstrated that disrupting base-pairing abrogates P-body localization, supporting a potentially two-step regulatory mechanism: (1) initial target recognition through sequence-specific base-pairing, followed by (2) P-body recruitment mediated by RBP interactions. The loss of repression upon *DDX6* or *LSM14A* knockdown underscores the functional necessity of P-body localization, aligning with its known role in translational silencing^50^.

Several lines of evidence underscore the biological specificity of this circRNA-mediated translational regulation: (1) Only circular *CDR1as* isoforms showed repressive activity, despite identical sequence to linear counterparts; (2) Mutations disrupting base-pairing abolished repression without affecting mRNA stability; (3) Effects were consistent across multiple cell types and endogenous targets. This specificity is unlike the proposed “sponge” functions, where circRNAs often show minimal effects on miRNA targets^63^. Our transcriptome-wide analyses further suggest this mechanism may be generalizable, with thousands of circRNAs showing P-body localization and target enrichment.

Previous studies have shown that *Cdr1as* KO mice exhibit impaired excitatory synaptic transmission with increased spontaneous vesicle release and a phenotype associated with neuropsychiatric disorders, namely a severe sensorimotor gating deficit^27^. As a target mRNA of *CDR1as*, *GRIN2B*—a subunit of N-Methyl-D-aspartate (NMDA) receptors^64^ has been reported as aberrantly upregulated in the anterior cingulate cortex of patients with major depression and the blood of epilepsy patients^65-^_67_. Consistent with these findings, we observed a significant upregulation of *GRIN2B* translation in *CDR1as* knockdown neurons. *MYO5A*, another mRNA target of *CDR1as*, has been shown to function in the transport and secretion of large dense-core vesicles (LDCVs) carrying neuronal peptides^68^. Its overexpression has been shown to impair LDCV movement and secretion in hippocampal neurons^69^. In this study, we also observed elevated *MYO5A* translation in *CDR1as* knockdown neurons. These findings suggest that upregulations of *GRIN2B* and *MYO5A* may contribute to the abnormal synaptic signaling transmission observed in *CDR1as* KO mice.

In summary, we describe a conserved pathway of translational regulation combining RNA-RNA base-pairing specificity with P-body-mediated silencing. This expands the functional repertoire of circRNAs while providing a framework for understanding how their unique biogenesis enables distinct regulatory capabilities compared to linear RNAs. Given the growing recognition of translational dysregulation in neurological disorders^70^, these findings may open new therapeutic avenues targeting circRNA-mRNA interactions. Additionally, our study illustrates a computational analysis strategy applicable to all RNA-RNA interaction datasets and provides a valuable data resource for the functional exploration of circRNA-target RNA interactions in various biological processes and diseases. Moreover, we developed a GRIC-seq method for global mapping of RNA-RNA spatial interactions in P-bodies, and this method should be generally applicable for studying RNA-RNA interactions in other membraneless granules.

### Limitations of the study

While our study demonstrates that circRNAs—but not their linear isoforms—repress target mRNA translation, the underlying mechanism remains incompletely understood. It is still unclear whether the circular topology primarily stabilizes base-pairing with target RNAs or facilitates specific interactions with P-body components to mediate translational repression. Although we established the necessity of P-bodies through the knockdown of core components DDX6 and LSM14A, the relative contributions of individual P-body proteins remain unresolved. Furthermore, whether circRNAs actively recruit silencing machinery or passively accumulate within these structures is an open question.

We also cannot exclude the possible involvement of direct ribosome inhibition mechanisms, such as steric hindrance or RNA structural rearrangements, which warrant further investigation. From a physiological perspective, circRNA-mediated translational control may be particularly important in neurons, where localized protein synthesis critically supports synaptic plasticity^71^. Future studies manipulating endogenous circRNA-target interactions in animal models will help establish their physiological relevance. Finally, RIC-seq data revealed previously uncharacterized interactions between circRNAs and noncoding exons, suggesting additional regulatory functions that remain to be explored.

## ACKNOWLEDGMENTS

We thank Drs. Changchang Cao and Bing Zhou for their help with data analysis; Dr. Jinyang Zhang from Fangqing Zhao’s lab for the discussion on circRNA identification and quantification. Dr. Xi Zhou for kindly providing *AGO2*-KO 293T and *DICER*-KO 293T cell lines; Dr. Yu Zhou for GFP-LSM14A and GFP-LSM14A-Δ cell lines; and Dr. Bing Zhu for the gifted HDAC2 antibody. We also thank the National Health and Disease Human Brain Tissue Resource Center for providing hippocampal tissues. This work was supported by the National Key Research and Development Program of China (2022YFA1303300), the National Natural Science Foundation of China (32025008, 32130064, and 81921003), and the Strategic Priority Program of Chinese Academy of Sciences (XDB37000000) to Y.X.; the Guangdong Major Project of Basic and Applied Basic Research (2023B0303000005) and Guangdong Provincial Special Support Program for Prominent Talents (2021JC06Y656) to P.Z.; the National Natural Science Foundation of China (32300457) to P.L.; the National Natural Science Foundation of China (32470610) to H.Z; and the China Postdoctoral Science Foundation (2024M763462 and GZB20240799) to J.L.

## AUTHOR CONTRIBUTIONS

Y.X. conceived and supervised the project. P.L. and Z.C. constructed the RIC-seq library of human and mouse hippocampal tissues, hNPC-derived neurons, and HT29 cells. H.Z. conducted the bioinformatics analysis with the help of Z.Y., H.L., and Y.Z. under the guidance of P.Z. R.Y. and B.L. established the KO and KI cell lines. R.Y. constructed the *CDR1as* knockdown pTRIPZ plasmids. P.L. performed RNA FISH, IF, hNPC differentiation, plasmid construction, ChIRP-MS, and luciferase assays. P.L. and J.L. performed RNA HCR-FISH. P.L. and Z.C. purified P-bodies for RIC-seq analysis. P.L. and H.Z. prepared the figures. X.L. provided the human hippocampal tissues. X.W. guided manuscript and figure preparations. Y.X. wrote the manuscript with contributions from P.L. and H.Z.

## SUPPLEMENTARY TABLES

Table S1. List of mapping results of RIC-seq datasets generated in the hippocampus and ten cell lines.

Table S2. List of circRNA and target RNA interactions supported by BSJ.

Table S3. List of circRNA and target RNA interactions supported by non-BSJ.

Table S4. List of DEGs detected by RNA-seq and Ribo-lite in hNPC-derived neurons upon *CDR1as* knockdown.

Table S5. List of *CDR1as*-associated proteins identified by ChIRP-MS in hNPC-derived neurons.

Table S6. List of circRNAs detected in purified P-bodies from HeLa cells.

Table S7. List of circRNAs detected in purified P-bodies from hNPC-derived neurons.

Table S8. List of PCR primers, shRNA, and siRNA used in this study.

Table S9. List of HCR-FISH, smFISH, and ChIRP probes used in this study.

### Data and code availability

High-throughput sequencing data generated in this study, including RIC-seq, GRIC-seq, RNA-seq, Ribo-lite, and total RNA-seq, have been deposited in the Genome Sequence Archive (GSA) under accession number CRA020865 and the Genome Sequence Archive for Human (GSA-Human) under accession number HRA009517. Custom codes used for data analysis can be found at https://github.com/zhanghm36/circTargetMap.

## METHODS

### Cell culture

The hNPCs were maintained on DMEM/F12 medium (10565018, Thermo Fisher Scientific) supplemented with 20 ng/ml FGF2 (233-FB-500, R&D), 0.5% N-2 (17502048, Thermo Fisher Scientific) and 1% B-27 (17504044, Thermo Fisher Scientific) in 10 μg/ml Pol-L-ornithine hydrobromide (P3655, Sigma-Aldrich) and 2.5 μg/ml laminin (23017015, Thermo Fisher Scientific) coated plates. The neuronal cells were differentiated from hNPCs using a previously established protocol^35,36^. Specifically, hNPCs were treated with 10 μM Y27632 (1245, Tocris) in differentiation medium (DMEM/F12 supplied with N-2 and B-27) for 1 day and replaced with differentiation medium every 2 days for 3 weeks. The 293T, *AGO2*-KO 293T, and *DICER*-KO 293T cells were provided by Dr. Xi Zhou at Wuhan Institute of Virology and maintained as previously described^72^. GFP-LSM14A and GFP-LSM14A-Δ HeLa cells were kindly provided by Dr. Yu Zhou at Wuhan University and maintained as previously described^54^. Lenti-X 293T (Takara, 632180) and HeLa cells (ATCC, CCL-2) were maintained on DMEM medium (C11965500BT, Thermo Fisher Scientific) supplemented with 10% FBS and 100 U/ml penicillin-streptomycin (15140, Life Technologies) at 37°C in 5% CO_2_. HT29 cells (ATCC, HTB-38) were cultured with McCoy’s 5A medium (16600082, Thermo Fisher Scientific) containing 10% FBS and 100 U/ml penicillin-streptomycin (Life Technologies). All cells were confirmed to be mycoplasma-free using PCR-based methods. The primers utilized for the mycoplasma test are listed in Table S8.

### Mice

The hippocampal tissues were isolated from the 10-week-old C57BL/6J mice (SPF Biotechnology Co., Ltd., China). All mice were maintained under specific pathogen-free conditions at 25°C. All animals were humanely euthanized by inhalation of carbon dioxide. The animal study protocol was approved by the Ethics Committee of the Institute of Biophysics, Chinese Academy of Sciences (SYXK2023188).

### Human hippocampus

Human hippocampus tissues were obtained from four donor patients at the Chinese Human Brain Bank, Zhejiang University School of Medicine. The donors signed informed consent. The collection and research plans for human tissues were approved by the Ethics Committee of Zhejiang University School of Medicine (#2020-005).

### Identification of circRNA-target RNA interactions by circTargetMap from RIC-seq data

We constructed RIC-seq libraries for HT29, HEK293T, hNPC-derived neurons, and hippocampal tissues of human and mouse as previously described^32,33^. The RIC-seq data of GM12878, H1-hESC, HeLa, HepG2, hNPC, IMR90, and K562 cell lines were downloaded from our previous works^32,37^. Adapters from paired-end sequencing data of RIC-seq libraries were first trimmed using Trimmomatic v.0.36 ^73^. PCR duplicates were removed with in-house scripts, as we previously described^32^. Poly(N) tails at the 3′ end were clipped using Cutadapt v4.1 ^74^. After filtering, the paired reads were aligned to the human or mouse pre-rRNA sequences. The remaining reads were then mapped to the human (hg19) or mouse (mm10) genomes using HISAT2 v2-2.1.0 ^75^ with default parameters to remove the normally mapped reads, including those spanning RNA splice junctions. The unmapped reads from read 1 and read 2 generated by HISAT2 were separately re-mapped to the reference genome using BWA v0.7.17 ^76^ with the following parameters: bwa mem -k 12 -T 15. Given the co-expression of circRNAs and their cognate linear RNAs in cells, we first identified the circRNA-target RNA interactions using circTargetMap (developed in this study, https://github.com/zhanghm36/circTargetMap) by analyzing chimeric reads with one arm mapped to the circRNA BSJ site and the other mapped to the target RNA. These interactions were termed as BSJ-supporting interactions. To comprehensively map target RNAs for the highly circularized circRNAs (circular-to-linear ratio ≥ 0.85), we expanded our analysis to include chimeric reads with one arm aligned to any region of the circRNA transcript and the other to the target RNA. These interactions were classified as non-BSJ-supporting interactions. All aligned read arms were then extracted and re-mapped to the genome using HISAT2 (v2.1.0) ^75^ and Bowtie2 ^77^, and only uniquely mapped arms (MAPQ > 20) were retained for interaction assignment. To distinguish biologically significant interactions from background noise, we implemented a Monte Carlo simulation^38^, comparing observed interactions with simulated random interactions through 100,000 iterations. Interactions with a *p*-value below 0.05 were considered statistically significant. For highly circularized circRNAs, we applied additional stringent criteria: only interactions supported by at least two chimeric reads and with a *p*-value below 0.05 were classified as high confidence.

### CircRNA reference resource

Human and mouse circRNAs were obtained from the circBase^5^ and circAtlas^6^ databases. We included all circRNAs from circBase, while only high-abundance circRNAs from circAtlas (CPM > 0.1 in at least one sample) were retained. Redundant circRNAs between the two databases were filtered based on the BSJ site and the spliced circRNA length. In cases of redundancy, circBase entries were prioritized.

### Identification of highly circularized circRNAs

To identify highly circularized circRNAs, we analyzed total RNA-seq data from multiple resources. Public datasets for human hippocampus (PRJNA322318)^78^ and seven cell lines were downloaded from the NCBI SRA database, including GM12878 (GSE78550)^79^, H1-hESC (GSE86189)^80^, HEK293T (GSE272077)^81^, HepG2 (GSE90229)^79^, HT29 (GSE78684)^79^, hNPC (PRJNA596331), and IMR90 (GSE90257)^79^. Additionally, we generated in-house total RNA-seq data for K562, HeLa, hNPC-derived neurons, and mouse hippocampus. CIRIquant^82^ was employed for circRNA characterization and quantification with default parameters. Briefly, reads were first aligned to either the human (hg19) or mouse (mm10) genome using BWA^83^. Candidate circRNAs were identified using CIRI2 ^84^. A pseudo-reference of the circular sequence is generated using the back-spliced region. Then, all candidate reads are re-aligned to the pseudo-reference sequence using HISAT2 (v2.1.0)^75^, with concordantly mapped read pairs spanning 10 bp region of the junction site are considered as circular reads. Non-circular reads aligned across the back-spliced junction sites are used to calculate the circRNA junction ratio. Only circRNAs with read count ≥5, circular-to-linear ratio ≥ 0.85, and detected in at least two replicates were considered as highly circularized circRNAs.

### MFE calculation for interacting circRNA and target RNA fragments

To assess the base pairing ability between circRNA and target RNA, we extracted 50-nt fragments upstream and downstream of the chimeric read junctions. The MFE for potential hybrids formed between these fragments was computed using the DuplexFold program from RNAstructure^39^ software with default parameters. Additionally, MFE values for shuffled sequences with identical nucleotide compositions were calculated (repeated 10 times) and served as negative controls. We applied the same method to analyze specific interactions involving those between *CDR1as* (and its compensatory mutants) and WT or mutant fragments of its non-BSJ targets (*GRIN2B*, *MYO5A*, and *AKAP9*), as well as between *circRMST* (and its compensatory mutants) with WT or mutant fragments of its BSJ targets (*APC2* and *PALS2*).

### Quantification of circRNA and target expression levels

To quantify circRNA and target RNA expression levels, we performed rRNA-depleted RNA sequencing for hNPC-derived neurons. For the human hippocampus, we obtained the published rRNA-depleted RNA-seq data from Duarte et al^85^. CircRNA abundance was quantified as BSJ read counts per million (CPM) using CIRIquant^82^, while target RNAs were measured in FPKM (Fragments Per Kilobase of transcript per Million mapped reads) via StringTie^86^. To enable cross-comparison, we present a conversion method by accounting for differences in quantification units and read length, ultimately expressing all measurements in CPM.

First, we define a normalization factor (N) to standardize read counts per kilobase (kb) of transcript length. Given that Illumina sequencing typically produces 150-nt reads, we compute the normalization factor as:

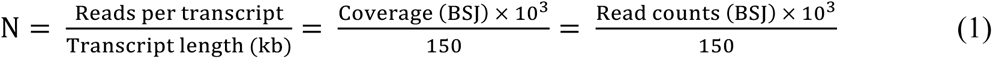

Here, CPM for circRNAs is calculated as:

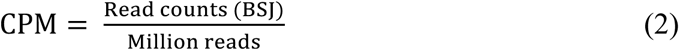

Meanwhile, FPKM incorporates transcript length normalization:

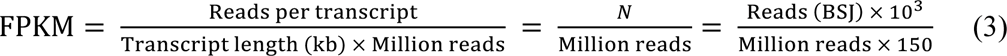

Substituting CPM into the equation yields the relationship:

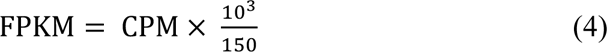

Conversely, CPM can be derived from FPKM as:

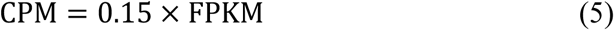

### ShRNA and siRNA knockdown

To knock down circRNAs, two shRNAs targeting the BSJ of *CDR1as* (sh*CDR1as*-1, sh*CDR1as*-2), *circRMST* (sh*circRMST*-1, sh*circRMST*-2), or *circMAN1A2* (sh*circMAN1A2*-1, sh*circMAN1A2*-2) were inserted into pLKO.1-EGFP-Puro plasmid between *EcoRI* and *AgeI* restriction sites, with the scrambled sequence (shScramble) used as a negative control. To inducibly knock down *CDR1as* or *circRMST* in hNPC-derived neurons, the shRNA sequence of *CDR1as* or *circRMST* was inserted into the pTRIPZ DOX-inducible shRNA vector between the *EcoRI* and *XhoI* restriction sites. As a negative control, the scramble sequence was inserted into the same backbone for comparison. For lentivirus packaging, pLKO.1-EGFP-Puro or pTRIPZ backbone-related constructs were mixed with psPAX2 and pMD2.G in a 4:3:1 ratio and transfected into Lenti-X 293T cells using Lipofectamine 2000 (11668019, Invitrogen) according to the manufacturer’s instructions. After 48 h and 72 h, the lentivirus-containing culture medium was collected, filtered through a 0.45 μm filter, and concentrated by centrifugation at 72,000 *g* in a SW28 rotor for 2 h at 4°C using an ultracentrifuge (Beckman Coulter). After removing the supernatant, the concentrated lentivirus was resuspended in PBS and stored at -80°C. For pLKO.1-EGFP-Puro lentivirus infections, two-week differentiated hNPC cells were infected with shRNAs or scrambled control containing pLKO.1-EGFP-Puro lentivirus (MOI = 10) supplemented with 1 μg/ml polybrene (TR-1003, Sigma-Aldrich) for 48 h. The cells were then selected with 0.25 μg/ml puromycin (A1113803, Thermo Fisher Scientific) for 72 h and replaced with differentiation medium for another 48 h. For pTRIPZ lentivirus infections, hNPCs were infected with *CDR1as* shRNA*, cricRMST* shRNA, or scramble shRNA containing lentivirus (MOI = 10) supplemented with 1 μg/ml polybrene for 48 h. The cells were then selected with 1 μg/ml puromycin for 72 h and differentiated for 2 weeks. Inducible knockdown was achieved by supplying 1 μg/ml Dox (631311, Clontech) to the culture medium for 7 days to induce the expression of RFP and knock down the expression of circRNAs in neurons.

For siRNA knockdown, we performed transfection with 40 pmol of previously validated siRNA targeting *DDX6*^87^, *LSM14A*^88^*, IGF2BP*1^89^ using Lipofectamine 2000. After 48 hours, cells were collected for qPCR or immunoblotting analyses. A non-targeting siRNA was used as a negative control. All siRNAs were synthesized by Hippobio (Huzhou, China). The sequences of shRNA and siRNA are listed in Table S8.

### GRIC-seq

The GRIC-seq method was designed to identify RNA-RNA interactions in membraneless granules, such as P-bodies and stress granules. GRIC-seq steps include granule purification, granule capture using magnetic beads, formaldehyde cross-linking, permeabilization, pCp-biotin labeling, proximity ligation, RNA purification, RNA fragmentation, biotin-labeled RNA enrichment, and strand-specific library construction.

#### Fluorescence-labeled P-body purification by FAPS

P-body purification from GFP-LSM14A stably expressing HeLa cells was performed using FAPS as previously described^49,54^. GFP-LSM14A and GFP-LSM14A-Δ HeLa cells were cultured in 15 cm dishes until reaching 80-90% confluency. Cells were washed twice with PBS and scraped off in 2 ml ice-cold PBS. Cell pellets were resuspended in lysis buffer (50 mM Tris, pH 7.4, 1 mM EDTA, 150 mM NaCl, 0.2% Triton X-100, and 1× EDTA-free protease inhibitor cocktail (20124ES, Yeasen Biotechnology) in the presence of 40 U/ml Ribolock RNase Inhibitor (EO0381, Thermo Fisher Scientific)) and incubated on ice for 5 min and then passed through a 1 ml syringe needle eight times. Nuclei and cell debris were removed by centrifugation at 1,000 *g* for 5 min at 4°C. 5% of the supernatant was saved as a cytoplasmic input control. Cytoplasmic and nuclear fractions were stained with Hoechst 33342 (5 μg/ml) at 37°C for 10 min. Hoechst 33342 fluorescence was analyzed by flow cytometry (Influx, BD) using a 405 nm excitation laser to verify the separation of nuclear and cytoplasmic fractions. The remaining supernatant was centrifuged at 10,000 *g* for 10 min. The resulting pellet was resuspended in 200 μl lysis buffer (defined as the pre-sorted fraction). The pre-sorted fraction was diluted in 2 ml lysis buffer and transferred to a FACS tube. Sorting was performed using a flow cytometer equipped with a 70 μm nozzle at 30 psi using 488 nm excitation laser (Influx, BD). GFP-LSM14A-Δ samples were used as a negative control. The sorting rate was maintained at approximately 5000 events per second for optimal accuracy. For P-body GRIC-seq, purified P-bodies were incubated with GFP-Trap^®^ beads (gtma-20, Chromotek) for 1 h at 4°C with rotation at 20 rpm. After incubation, the samples were washed three times with dilution buffer (10 mM Tris-HCl, pH 7.4, 0.5 mM EDTA, 150 mM NaCl). The purified P-bodies were subsequently crosslinked for GRIC-seq library construction.

#### Non-fluorescently-labeled P-body purification by immunoprecipitation

For non-fluorescently labeled P-bodies in neuronal samples, we isolated them through antibody-based immunoprecipitation following established protocols^55,56^. For each IP, 50 µl of Protein A/G magnetic beads (88803, Thermo Fisher Scientific) were washed three times with 0.1 M sodium phosphate buffer (0.05% Tween-20, pH 8.0) and then resuspended in the same buffer containing 20 µg of anti-LSM14A antibody (sc-398552, Santa Cruz Biotechnology). After incubation for 1 h at 25°C on a rotator at 20 rpm, the beads were washed three times with 600 µl of wash buffer (1× PBS, 0.1% SDS, 0.5% sodium deoxycholate, 0.5% NP-40). The cytoplasmic supernatant from neurons was incubated with anti-LSM14A-coupled beads for 2 h at 4°C with rotation at 20 rpm to pull down P-bodies. The pulled P-bodies were washed four times with wash buffer and saved for GRIC-seq library construction.

#### Cross-linking and Permeabilization

P-body coupled magnetic beads were resuspended with 1 ml PBST (0.01% Tween 20). To fix the protein-mediated RNA-RNA interactions in P-bodies, 27 µl of 37% (m/v) formaldehyde was added to the sample and incubated at 25°C for 10 min. The reaction was quenched by adding 1/20 volume of 2.5 M glycine and incubated at 25°C for 10 min. After washed three times with PBST, the magnetic beads were resuspended in 1 ml of permeabilization buffer (10 mM Tris-HCl pH 7.5, 10 mM NaCl, 0.5% NP-40, 0.3% Triton X-100, 0.1% Tween 20, 1× protease inhibitors (Yeasen Biotechnology), 2 U/ml SUPERase In RNase inhibitor (Thermo Fisher Scientific) and incubated on ice for 15 min with mixing every 2 min. Then the beads were washed three times with 1× PNK buffer (50 mM Tris-HCl pH 7.4, 10 mM MgCl_2_, 0.1% Tween 20).

#### MNase treatment and pCp-biotin labeling

The thoroughly washed beads were resuspended in 200 μl of 1 × MN mixture (50 mM Tris-HCl pH 8.0, 5 mM CaCl_2_, 0.03 U/μl micrococcal nuclease (EN0181, Thermo Fisher Scientific)), and incubated at 37°C for 10 min. After being washed twice with 1× PNK + EGTA buffer (50 mM Tris-HCl pH 7.4, 20 mM EGTA, 0.1% Tween 20) and twice with 1× PNK buffer, the beads were resuspended in 100 μl of 1 × Fast AP reaction buffer with 10 U of FastAP alkaline phosphatase (EF0651, Thermo Fisher Scientific) and incubated for 15 min at 37°C. The reaction was stopped by sequentially washing twice with 1× PNK + EGTA buffer, twice with 1× high-salt wash buffer (5× PBS, 0.1% Tween 20), and three times with 1× PNK buffer. To label RNA 3’ end with pCp-biotin, the beads were resuspended in a ligation mixture containing 10 μl of 10 × RNA ligase reaction buffer, 6 μl of 40 U/μl RNase inhibitor, 4 μl of 1 mM pCp-biotin (20160, Thermo Fisher Scientific), 10 μl of 10 U/μl T4 RNA ligase (EL0021, Thermo Fisher Scientific), 20 μl of nuclease-free water, and 50 μl of 30% PEG 20000. The sample was incubated at 16°C overnight.

#### Proximity ligation

After pCp-biotin labeling, the beads were washed three times with 1× PNK buffer and resuspended in a PNK mixture containing 10 μl of 10 × Imidazole buffer (500 mM imidazole-HCl pH 6.4, 100 mM MgCl_2_), 6 μl of 10 mM ATP, 4 μl of 10 U/μl T4 polynucleotide kinase (EK0032, Thermo Fisher Scientific), 10 μl of 0.1 M DTT, 70 μl of nuclease-free water, and incubated for 45 min at 37°C. For in situ proximity ligation the beads were resuspended in a ligation mixture containing 20 μl of 10× RNA ligase reaction buffer, 20 μl of 1 mg/ml BSA, 8 μl of 40 U/μl RNA inhibitor (Thermo Fisher Scientific), 10 μl of 10 U/μl T4 RNA ligase (Thermo Fisher Scientific), 142 μl of nuclease-free water, and incubated at 16°C overnight.

#### RNA purification and library construction

On the next day, the beads were washed three times with 1× PNK buffer, then resuspended in 220 μl of proteinase K buffer (10 mM Tris-HCl pH 7.5, 10 mM EDTA, 0.5% SDS) and 30 μl of proteinase K (9034, Takara). The sample was incubated at 37°C for 60 min and 56°C for 15 min. After flash centrifugation, the tube was placed on a magnet stand for 1 min. The supernatant was transferred to a new tube and mixed with 750 μl of TRIzol LS (10296028, Thermo Fisher Scientific). After adding 200 μl of chloroform and vortexing for 15 s, the tube was centrifuged at 16,000 *g* for 15 min at 4°C. The supernatant was transferred to a new 1.5 ml Eppendorf tube and mixed with 500 μl of isopropanol and 1 μl of glycoblue (15 mg/ml, AM9515, Thermo Fisher Scientific). After overnight incubation at -20°C, the RNA was pelleted at 16,000 g for 20 min at 4°C. The RNA pellet was resuspended in 10 μl of nuclease-free water after washing twice with 75% ethanol. The subsequent steps of GRIC-seq were performed exactly as we previously described^32^, including RNA fragmentation, pCp-biotin selection, and strand-specific library preparation.

### Overexpression of circRNAs in hNPC-derived neurons

The full-length sequences of *CDR1as* or *circRMST* were amplified from neuronal cDNAs and cloned into the circRNA overexpression vector pLO5-ciR (GENESEED) between the *EcoRI* and *BamHI* restriction sites. As a control, a fragment from λDNA matching the length of *CDR1as* or *circRMST* was inserted into the same vector. Lentiviral packaging, infection, and puromycin selection were performed as described above.

### RNA extraction, reverse transcription, and qPCR

Total RNAs were extracted from cells with TRIzol reagent (15596026, Thermo Fisher Scientific) following the manufacturer’s instructions. For reverse transcription, 1 μg of total RNA was first treated with RQ1 RNase-free DNase (M6101, Promega) to remove potential DNA contamination. Subsequently, reverse transcription was performed using M-MLV reverse transcriptase (M1701, Promega) and random hexamers. The cDNA was diluted (1:5) and served as templates for qPCR analysis using 2X Hieff qPCR master mix (11203ES08, Yeasen) and specific primers. The qPCR primers used in this study are listed in Table S8.

### Immunoblotting

Cells were lysed using RIPA lysis buffer (89901, Thermo Fisher Scientific), and the protein concentrations were quantified using BCA Protein Assay Kits (23225, Thermo Fisher Scientific) following the manufacturer’s instructions. Immunoblotting was performed as previously described^90^. The anti-MAN2A1 (1:1000, CSB-PA613697LA01HU, CUSABIO), anti-MTURN (1:1000, 27936-1-AP, Proteintech), GRIN2B (1:1000, 66565-1-Ig, Proteintech), anti-MYO5A (1:1000, YT2950, Immunoway), anti-AKAP9 (1:1000, CSB-PA857882LA01HU, CUSABIO), anti-APC2 (1:1000, sc-517022, Santa Cruz), anti-PALS2 (1:000, ab180508, Abcam), anti-STAT3 (1:2000, ab68153, Abcam), anti-HDAC2 (1:2000, ab32117, Abcam), anti-ZFP36L1 (1:1000, ab230507, Abcam), anti-PTPRM (1:1000, YN2112, Immunoway), anti-MAP3K2 (1:2000, ab33918, Abcam), anti-POLR2C (1:1000, AP20520b, INSIGHT BIOTECHNOLOGY), anti-TUBB (1:10000, 66240-1-Ig, Proteintech), anti-AGO2 (1:2000, ab186733, Abcam), anti-DICER (1:1000, Ab259327, Abcam), anti-ACTB (1:5000, AC026, Abclonal), anti-HNRNPU (1:500, sc-32315, Santa Cruz), anti-IGF2BP1 (1:1000, RN007P, MBL), anti-DDX6 (1:2000, ab307418, Abcam), anti-LSM14A (Santa Cruz Biotechnology), HRP-coupled goat anti-mouse IgG (H+L) (1:5000, 31430, Invitrogen), HRP-coupled goat anti-rabbit IgG (H+L) (1:5000, 31460, Invitrogen) antibodies were used in this study for immunoblotting.

### Immunofluorescence

The cells on coverslips were fixed with 3.7% formaldehyde for 10 min at 25°C and washed three times with PBS. Cells were then permeabilized with 0.5% Triton X-100 for 20 min at 25°C and washed three times with PBS. Subsequently, the cells were blocked with 5% BSA (B2064, Sigma-Aldrich) for 1 h at 25°C, followed by overnight incubation with primary antibodies in 1% BSA at 4°C. After washing three times with PBS, the cells were incubated with fluorescently labeled secondary antibodies in the dark at 25°C for 1 h. Following incubation, cells were washed three times with PBS. Finally, the stained cells were mounted with a DAPI-contained mounting medium (H-1200, VECTOR) following the manufacturer’s instructions. The anti-MAP2 (1:200, MAB3418, Millipore), anti-TUJ1 (1:200, 802001, BioLegend), anti-DDX6 (1:200, ab307418, Abcam), Alexa Fluor™ 594 labeled donkey anti-mouse IgG (1:200, A21203, Invitrogen), and Alexa Fluor™ 488 labeled donkey anti-rabbit IgG (1:200, A21206, Invitrogen) antibodies were used for IF analysis. Images of stained cells were captured using a Nikon-Ti Confocal laser scanning microscope.

### HCR-FISH

HCR-FISH of *CDR1as*, *MAN2A1*, *MTURN*, *FBH1*, *RARS2*, *POLR2C, circRMST, APC2, PALS2, CYB5R4, circMAN1A2, PTPRM, MAP3K2* were performed as previously described^40^. Cells were fixed with 3.7% formaldehyde for 10 min at 25°C, followed by washing twice with PBS. Subsequently, cells were permeabilized with 70% ethanol (vol./vol.) overnight at -20°C. Then, cells were prehybridized with HCR hybridization buffer (5× SSC, 9 mM citric acid, 50 μg/ml heparin, 1× Denhardt’s solution, 0.1% Tween-20, 30% formamide, and 10% dextran sulfate) for 30 min at 37°C. After twice washing with 2× SSC, cells were incubated with probes (2 pmol for each RNA) in HCR hybridization buffer at 37°C overnight. The sections were washed four times for 5 min with wash buffer (5× SSC, 9 mM citric acid, 50 μg/ml heparin, 1× Denhardt’s solution, 0.1% Tween-20, and 30% formamide). The signals in cells were amplified by incubating with 18 pmol denatured hairpin probes at room temperature overnight, followed by five times washes with 5× SSCT buffer (5× SSC, 0.1% Tween-20) in the dark. Finally, the stained cells were mounted with a DAPI containing mounting medium (VECTOR). For combined IF and HCR-FISH, cells were first processed for IF as described above. After IF staining, cells were re-fixed with 3.7% formaldehyde. HCR-FISH was then performed as previously described, except that the permeabilization step was omitted. HCR-FISH probes used in this research are listed in Table S9. A Nikon-Ti Confocal laser scanning microscope was used to capture the stained cell images.

### smFISH

The FISH probes were designed using Stellaris probe designer (https://www.biosearchtech.com/support/tools/design-software/stellaris-probe-designer) and synthesized by DIA-UP BIOTECH with Quasar 570 or 670 dye. smFISH analysis of *CDR1as*, GRIN2B, *MYO5A*, *AKAP9* and *POLR2C* were performed as previously described^32^. Briefly, cells were fixed with 3.7% formaldehyde for 10 min at 25°C, followed by two washes with PBS. Subsequently, cells were permeabilized with 70% ethanol (vol./vol.) for at least 1 h at 4°C. After permeablization, cells were washed with the wash buffer (2× SSC, 10% deionized formamide) at 25°C for 5 min. Next, cells were incubated overnight with the probe mixture (12.5 pmol for each RNA) in a hybridization buffer in a dark chamber at 37°C. Following hybridization, cells underwent two washes with wash buffer at 37°C for 30 min in the dark. Finally, the stained cells were mounted with DAPI-containing mounting medium (VECTOR) for imaging. IF and smFISH co-staining was performed as previously described^91^. Cells were first processed for IF, as described above. After completing the IF staining, cells were re-fixed with 3.7% formaldehyde at 25°C for 10 min and washed three times with PBS. Subsequently, the smFISH procedure was performed as described earlier, omitting the permeabilization step. smFISH probes used in this research were listed in Table S9. A Nikon-Ti Confocal laser scanning microscope was used for capturing images of stained cells.

### RNA-seq

The rRNA-depleted and strand-specific RNA-seq libraries were prepared as described previously^32^. Total RNA from hNPC-derived neurons, mouse hippocampal tissues, and purified P-bodies was extracted using TRIzol Reagent, and rRNA was removed using the Ribo-off rRNA Depletion Kit (Vazyme, N406-01). Fragmented, rRNA-depleted RNA was used for first-strand cDNA synthesis with random hexamer primers. Strand specificity was ensured by incorporating dUTP in the dNTP mix during second-strand cDNA synthesis. The cDNA was subsequently subjected to end-repair, dA-tailing, and adaptor ligation, followed by USER enzyme (NEB, M5505S) treatment and PCR amplification.

### Smart-seq2 library preparation

Smart-seq2 libraries were conducted as previously described^41^. Briefly, 10 ng of total RNA was used for reverse transcription and template switching, followed by pre-PCR using ISPCR primers. The amplified cDNAs were fragmented using Tn5 transposase with adaptor assembly (TD502, Vazyme), followed by PCR amplification for library preparation.

### Smart-seq2 data processing

Adaptors were first trimmed from paired-end sequencing reads by Trim Galore v0.4.2 (https://github.com/FelixKrueger/TrimGalore). The clean reads were subsequently mapped to the human (hg19) transcriptome by STAR v2.5.2b^92^ with the parameters: - outFilterMultimapNmax 1 -outSAMstrandField intronMotif. Gene expression levels were calculated by Cufflinks v2.2.1 ^93^ based on the hg19 refFlat annotations from the UCSC genome browser. The average FPKM from two replicates was calculated, and genes with FPKM >1 in at least one sample were considered to be expressed.

### Ribo-lite library preparation

The ribo-lite libraries were generated as previously described^42^. Briefly, cells were treated with 100 μg/ml cycloheximide (CHX, GC17198, GLPBIO)for 5 min at 25°C and then washed three times with CHX-containing PBS at 4°C. Next, cells were lysed by adding 300 μl lysis buffer (20 mM Tris pH 7.4, 150 mM NaCl, 5 mM MgCl_2_, 1 mM DTT, 1% Triton X-100, 100 μg/ml CHX) for 10 min, and spun at 20,000 *g* for 10 min at 4°C. The supernatant was treated with 1 μl RNase I (AM2295, Ambion) at 22°C for 45 min. The digestion was stopped by adding 10 μl SUPERase•In (AM2696, Ambion). Then, the supernatant was carefully layered onto a 1 M sucrose cushion prepared in lysis buffer. After centrifugation for 4 h at 260,000 *g* at 4°C, the ribosome-bound RNA from the pellet was extracted using TRIzol reagent. To recover RPFs, the extracted RNA was denatured and separated on a 15% (wt/vol) polyacrylamide TBE-urea gel via electrophoresis. The monosome RPFs region of 26 to 34 nucleotides was excised from the gel and purified. The purified RPFs were converted into libraries using the D-Plex Small RNA-seq Kit (Diagenode, C05030001) following the manufacturer’s instructions.

### Ribo-lite data processing

The UMI, TMI motif, and adaptors were trimmed with Cutadapt v4.1 ^74^ using the parameters: cutadapt --trim-n --match-read-wildcards -u 16 -n 3 -a AGATCGGAAGA GCACACGTCTG -a AAAAAAAA -m 15. The trimmed reads were first aligned to human rRNA sequences using Bowtie2 v2.3.4.3 ^77^ with the parameters: -seedlen = 23 to remove rRNA. The remaining reads were subsequently mapped to the transcriptome of human (hg19) using STAR v2.5.2b ^92^ with the following parameters: - outFilterMismatchNmax 2 -outFilterMultimapNmax 1 -outFilterMatchNmin 16 - alignEndsType EndToEnd. Gene expression levels were calculated using Cufflinks v2.2.1 ^93^ based on the CDS region defined by the hg19 refFlat annotations from the UCSC genome browser. The average FPKM for replicates was calculated, and genes with FPKM >1 in at least one sample were considered to be expressed.

### Identification of DEGs

DEGs were identified from RNA-seq and Ribo-lite data using DESeq2 ^94^. Gene read counts were quantified using featureCounts v2.0.3 ^95^ for both RNA-seq and Ribo-lite datasets. These counts were then analyzed by DESeq2 for differential expression analysis. Genes were considered differentially expressed if they met the following thresholds: |fold change| > 1.5, *p*-value < 0.05, and FPKM > 1 in at least one condition.

### TE analysis

TE was calculated as the ratio of Ribo-lite to RNA-seq (FPKM + 1/FPKM + 1). Differential TE analysis was conducted using DESeq2 ^94^, with TE > 1.5 and *p*-value < 0.05 classified as high TE, and TE < 0.67 and *p*-value < 0.05 classified as low TE.

### Dule luciferase assay

The fragments of *GRIN2B* 3’UTR, *MYO5A* 3’UTR, *APC2* 3’UTR, *PALS2* 3’UTR, and *AKAP9 CDS*, along with their respective mutant fragments and the corresponding compensatory mutant fragments of *CDR1as* and *circRMST*, were synthesized by Tsingke (Beijing). The four 3’UTR fragments and their mutants were inserted into the psiCHECK-2 backbone plasmid between the *XhoI* and *NotI* restriction sites. A fragment of γDNA was inserted into the same backbone as a control. For *AKAP9*, the CDS fragment and its synonymous mutant were homologously recombined into the psiCHECK-2 vector downstream of the “ATG” of the Renilla luciferase sequences. The compensatory mutants of *CDR1as* and c*ircRMST* were inserted into pLO5-ciR between the *EcoRI* and *BamHI* restriction sites. Additionally, the entire sequence of *CDR1as* or *circRMST* was inserted into pcDNA3 between *XhoI* and *BamHI* restriction sites for the overexpression of their linear isoform. The entire sequence of *CDR1as* or *circRMST* was recombined homologously into a forward circular frame-deficient pLO5-ciR. For gradient overexpressing *CDR1as*, the pLO5-ciR plasmid containing *CDR1as* was transfected into 293T cells, *AGO2*-KO 293T cells, and *DICER*-KO 293T cells with varying amounts (0, 250, 500 ng) and supplied to 500 ng using γDNA fragment inserted plasmid in a 24-well plate with Lipofectamine 2000 for 24 h. For the dual-luciferase assay, either the circRNA overexpression plasmid, the linear-*circRNA* overexpression plasmid, or the forward circular frame-deficient circRNA overexpression plasmid, along with the psiCHECK2 vector containing either the target or control fragment, was co-transfected into 293T or HeLa cells in 24-well plates using Lipofectamine 2000. For the *IGF2BP1* rescue experiment, the full *IGF2BP1* CDS sequence was inserted into pcDNA3 between the XhoI and BamHI restriction sites for overexpression to rescue *IGF2BP1* knockdown in 293T cells. After 24 h of transfection, cells were washed three times with PBS and lysed with 100 μl of cell lysis buffer (E1910, Promega) for 10 min at 25°C. Half of the lysate was then centrifuged at 12,000 *g* for 10 min at 4°C, and 10 μl of the supernatant from each sample was used to measure Firefly and Renilla luciferase activities using the Dual-Luciferase Reporter Assay System (E1910, Promega) on a GloMax system (Promega), following the manufacturer’s instructions. The other half of the lysate was used to extract total RNA using TRIzol LS (Thermo Fisher Scientific). Subsequently, reverse transcription and qPCR of Firefly and Renilla luciferase genes were performed as previously described^91^.

### CRISPR-Cas9-mediated KO and knock-in

The sgRNAs were designed using the CRISPR design tool CRISPOR^96^, and BLAST analysis was performed to ensure each sgRNA targeted a single genomic region. For KO of *GRIN2B* and *MYO5A* fragments in the 3′UTR, two sgRNAs were designed to flank the target region**–**one upstream and one downstream. The upstream sgRNAs were cloned into the px330-mCherry plasmid, and the downstream sgRNAs were inserted into the px458-EGFP plasmid at the BpiI restriction site. The paired sgRNA plasmids were co-transfected into 293T cells. To establish synonymous codon knock-in mutations in *AKAP9*, a sgRNA targeting the knock-in region was inserted into the px330-mCherry plasmid, and the donor construct containing the synonymous mutations was cloned into the pEGFP-N1 plasmid at the *NotI* restriction site. The sgRNA and donor plasmids containing synonymous mutations were co-transfected into 293T cells. Double-positive cells (mCherry⁺/GFP⁺) were individually sorted for culture and genotyping. Successfully edited KO or knock-in cells were confirmed by Sanger sequencing and used for downstream analyses.

### ChIRP-MS

ChIRP-MS was performed as previously described with minor modifications^48^. hNPC-derived neurons were fixed with 3% formaldehyde for 30 min and then quenched with 0.125 M glycine for 5 min at 25°C. Cell pellets (100 mg) were dissolved in 1 ml of ChIRP cell lysis buffer (50 mM Tris–HCl pH 7.0, 10 mM EDTA, 1% SDS, 1 mM PMSF (329-98-6, Sigma-Aldrich), protease inhibitors) and sonicated on ice for 5 min using a parameter of 3s on, 10s off cycle at 40% amplitude. The lysate was centrifuged at 16,000 *g* for 10 min at 4°C, and 5% of the supernatant was saved as input. 30 μl washed Myone Streptavidin C-1 Dynabeads (65002, Invitrogen) were added to the supernatant and incubated at 37°C with rotation for 30 min for pre-clearing. For the ChIRP-MS procedure, the precleared supernatant was mixed with 2 volumes of ChIRP hybridization buffer (750 mM NaCl, 1% SDS, 50 mM Tris-Cl pH 7.0, 1 mM EDTA, 15% formamide, 1 mM PMSF, protease inhibitors), and 1 μl biotin-labeled probes mix (100 μM). The mixture was then incubated at 37°C with rotation for 12 h. After hybridization, 100 μl of washed C1 beads were added to the reaction and incubated at 37°C with rotation for another 30 min. The supernatant was removed after placing the tubes on a Dynamag magnet for 5 min, and the beads were washed 5 times with ChIRP wash buffer (2× SSC, 0.5% SDS, 1 mM PMSF). To quantify the relative RNA enrichment by qPCR, 5% of the beads were transferred to a fresh tube after the last wash for RNA extraction and reverse transcription. To release proteins from the remaining beads, 50 μl 1× LDS loading buffer (NP0007, Invitrogen) was added and incubated at 95°C for 10 min. The eluted proteins were separated by electrophoresis on a 4-12% (wt/vol) Bis-Tris gel (NP0321, Thermo Fisher Scientific) and visualized by silver staining kit (24612, Thermo Fisher Scientific) according to the manufacturer’s instructions. The eluted proteins were subjected to immunoblottings or liquid chromatography with tandem mass spectrometry (LC-MS/MS). The probes used in this study are listed in Table S9.

### Sample preparation and mass spectrometry

The protein bands of interest in the silver gel were excised for in-gel digestion and subsequently identified by mass spectrometry. Briefly, proteins were reduced with 25 mM DTT and alkylated with 55 mM iodoacetamide. In-gel digestion was performed overnight at 37°C using sequencing grade-modified trypsin in 50 mM ammonium bicarbonate. Peptides were extracted twice with 1% trifluoroacetic acid in a 50% acetonitrile aqueous solution for 30 min. The peptide extracts were then concentrated using a SpeedVac (Savant™ SpeedVac™ SPD120, Thermo Fisher Scientific). For LC-MS/MS analysis, peptides were separated by a 60 min gradient elution at a flow rate of 0.3 μl/min with a Thermo-Dionex Ultimate 3000 HPLC system, directly interfaced with a Thermo Orbitrap Fusion mass spectrometer. The analytical column was a homemade fused silica capillary column (75 μm ID, 150 mm length; Upchurch, Oak Harbor, WA) packed with C-18 resin (300 A, 5 μm; Varian, Lexington, MA). Mobile phase A consisted of 0.1% formic acid, and mobile phase B consisted of 100% acetonitrile and 0.1% formic acid. The Orbitrap Fusion mass spectrometer was operated in the data-dependent acquisition mode using Xcalibur 3.0 software. A single full-scan mass spectrum was performed in the Orbitrap (350-1550 m/z, 120,000 resolution), followed by 3s of data-dependent MS/MS scans in an Ion Routing Multipole at 30% normalized collision energy (HCD). The MS/MS spectra from each LC-MS/MS run were searched against the human UniProtKB database using the Proteome Discovery searching algorithm v1.4.

### LC-MS/MS data processing

Protein abundance quantified by Proteome Discoverer was log2-transformed and averaged across replicates. Differential protein enrichment between LacZ and *CDR1as* ChIRP was assessed using a two-tailed, unpaired Student’s t-test.

### Identification of P-body-associated circRNAs

We employed CIRIquant^82^ for circRNA characterization in the biochemically purified P-bodies from hNPC-derived neurons, as well as FAPS purified P-bodies from HeLa cells stably expressing GFP-LSM14A^54^. The sequencing reads from rRNA-depleted RNA-seq were first mapped to the human genome (hg19). CircRNAs were considered detectable when supported by at least one read spanning the back-splice junction in any biological replicates.

### RNA-seq data analysis

Using rRNA-depleted RNA-seq, we quantified circRNA target expression in neuronal cytoplasmic and P-body fractions, as well as in HeLa cell cytoplasmic and P-body fractions, as previously described^54^. The raw paired-end reads were first processed with Trim Galore v0.4.2 to remove adapters, and ribosomal reads were filtered out by Bowtie2 (v2.3.4.3)^77^. The remaining reads were then aligned to the human genome (hg19) using HISAT2 (v2.1.0)^75^, and only uniquely mapped reads were retained for gene expression quantification by StringTie (v2.1.6)^86^. P-body enrichment of individual RNA was defined as the ratio of their relative expression level in P-body RNA-seq to that in the cytoplasm RNA-seq.

### TE analysis of P-body circRNA targets

To assess the TE of circRNA target mRNAs localized in P-bodies of HeLa cells, we downloaded polysome profiling and cytoplasmic RNA-seq data from the study by Shan et al. (HRA002519)^54^. For circRNA target mRNAs detected in neuronal P-bodies, we used ribosome profiling and RNA-seq data from the study by Wang et al. (GSE156671)^97^. The quantification of gene expression followed the same procedure as RNA-seq and Ribo-lite. TE of individual RNA was defined as the ratio of (FPKM + 1) from polysome RNA to (FPKM + 1) from cytoplasmic RNA for HeLa cells, or (FPKM + 1) of ribosome profiling to (FPKM + 1) from RNA-seq for neurons.

### Pathological variants analysis

Disease-associated variants were obtained from the GWAS Catalog^57^ (accessed May 13, 2025), and 78,298 were retained after manual curation. An additional 251,116 pathogenic or likely pathogenic variants were extracted from the ClinVar database^58^. To assess the enrichment of disease-associated variants around circRNA-target RNA interaction sites, we analyzed ±500 bp flanking regions of each chimeric junction. Using bedtools^98^, we calculated variant density across these regions after partitioning them into 10 bp bins. Similar analyses were conducted on randomly shuffled genomic regions of matching size to serve as a negative control. To evaluate how these variants affect circRNA-target RNA interaction stability, we used RNAplex (ViennaRNA v2.4.18)^99^ to model WT and mutant duplex structures. MFE values were calculated for all interactions, and variants were considered as high impact if they met both of the following criteria: (1) | Δ MFE| ≥ 3 kcal/mol, and (2) ≥ 20% MFE change compared to the WT.

### GO biological process enrichment analysis

GO enrichment analysis was performed on target genes affected by disease-associated variants using the Metascape web tool. Processes with a false discovery rate (FDR) < 0.05 were considered significant. The networks between genes and enriched processes were visualized using Cytoscape v.3.8.2 ^100^.

### Quantification and statistical analysis

Band intensities for immunoblottings were quantified with ImageJ^101^. The GraphPad Prism software was used to prepare figures and perform statistical analysis for all the qPCR and relative luciferase activity results. Pearson’s correlation coefficient, calculated using the R function cor.test, was used to assess the reproducibility of the RNA-seq and Ribo-seq data shown in Figures S3A and S3B. Box plots in Figures S1F 6C, 6D, 6F, and 6G were generated by the Python function df.boxplot, with *p*-values determined by the unpaired two-tailed Wilcoxon rank-sum test. For the box plot figures, the center line represents the median, the box borders denote the first (Q1) and third (Q3) quartiles, and the whiskers are the most extreme data points within 1.5× the interquartile range (from Q1 to Q3).

## SUPPLEMENTARY FIGURES AND LEGENDS

**Figure S1.**
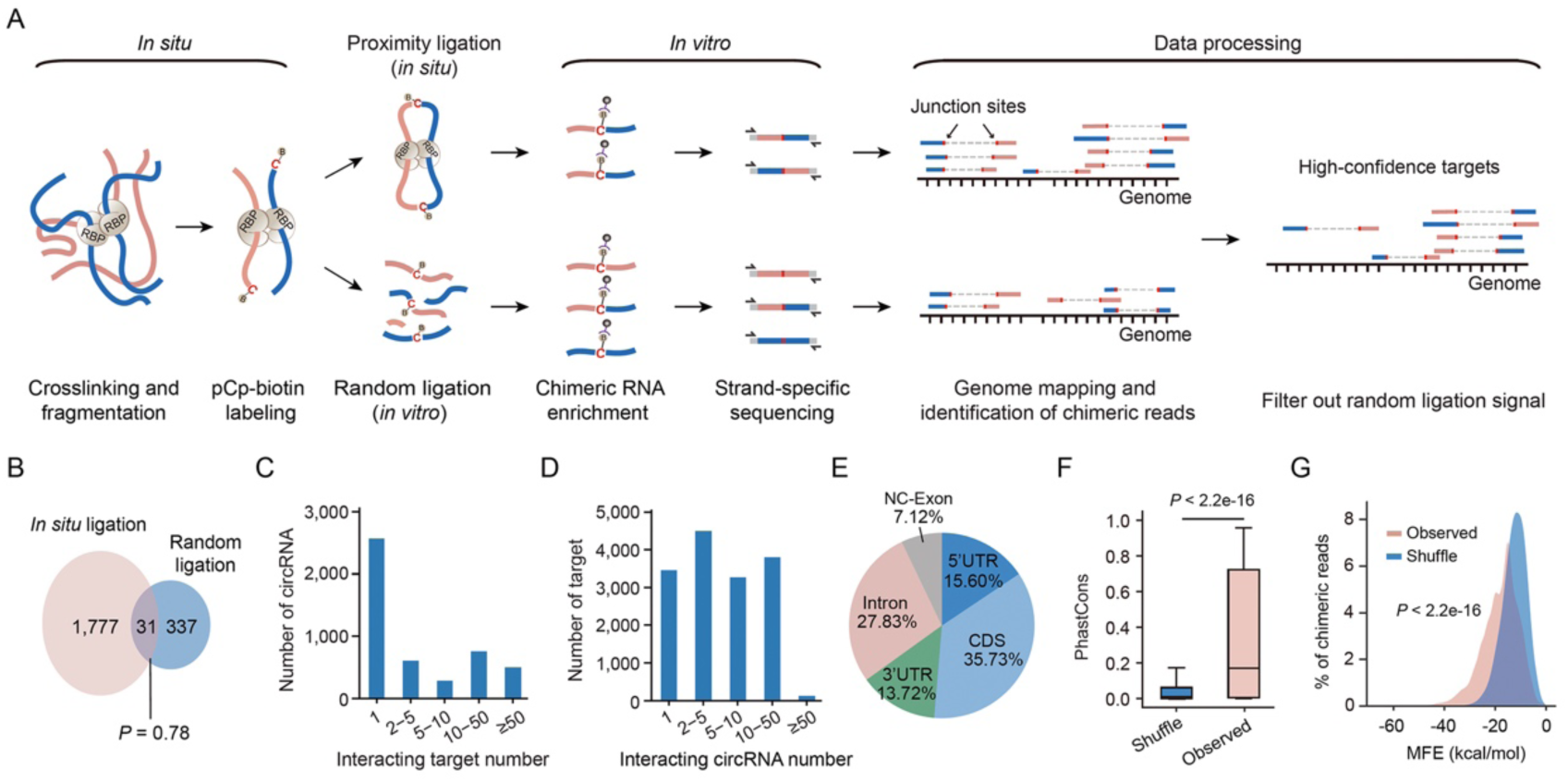
Comparison of *in situ* circRNA-target RNA proximity ligations with *in vitro* random ligations. (A) Diagram of the data analysis strategy for identifying high-confidence circRNA-target RNAs by excluding randomly ligated circRNA-target RNA interactions. (B) Venn diagram displaying the overlap of circRNA-target RNA interactions identified in proximity-ligated and random-ligated RNA fragments in hNPC-derived neurons. The *p*-value was calculated using a one-tailed hypergeometric test. (C) Bar plots showing the distribution of the interacting target RNA number of circRNAs. (D) Bar plots showing the distribution of the interacting circRNA number of target RNAs. (E) Pie chart showing the distribution of circRNAs interaction sites on target RNAs across the examined human samples. (F) Box plots showing the PhastCon scores in vertebrates for circRNA-interacting target RNA fragments compared to randomly shuffled sequences across the examined human samples. The *p*-value was calculated using a Wilcoxon rank-sum test. (G) Density plots displaying the MFE of base pairing between circRNA and target RNA fragments compared to randomly shuffled sequences across the examined human samples. The *p*-value was calculated using a Wilcoxon rank-sum test.

**Figure S2.**
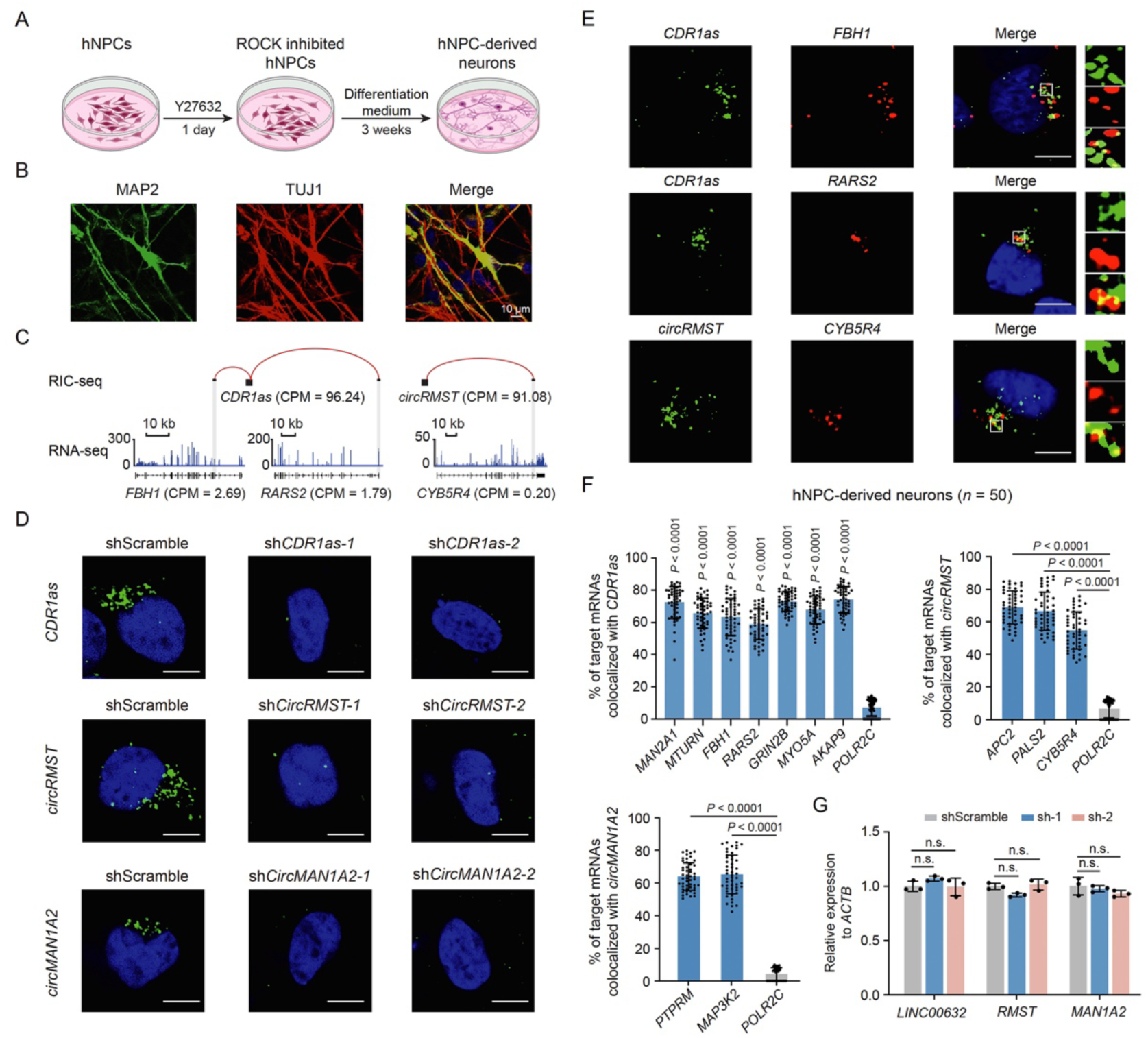
CircRNAs colocalize with target mRNAs in hNPC-derived neurons. (A) Diagram of the protocol for differentiating hNPCs into neurons. (B) IF results of neurons co-stained against MAP2 (green) and TUJ1 (red). Scale bar, 10 μm. (C) Snapshots of RIC-seq and RNA-seq tracks showing interactions of *CDR1as* and *circRMST* with their respective target mRNAs in neurons. (D) HCR-FISH detection of *CDR1as*, *circRMST*, and *circMAN1A2* in neurons treated with circRNA-specific shRNAs versus scramble shRNA. DAPI, blue; *circRNAs*, green. Scale bar, 5 μm. (E) HCR-FISH shows the colocalization of circRNAs with their target mRNAs in neurons. Target mRNAs, red. Scale bar, 5 μm. (F) Percentage of colocalization between circRNAs (*CDR1as*, *circRMST*, and *circMAN1A2*) and their target mRNAs (*n* = 50 neurons per group). *POLR2C* serves as a non-target control mRNA. Data are presented as mean ± s.d; two-tailed unpaired Student’s *t*-test. (G) qPCR analysis of linear transcript levels (*LINC00632*, *RMST*, and *MAN1A2*) in circRNA-depleted neurons versus shScramble-treated controls. Data are presented as mean ± s.d.; *n* = 3 independent replicates, two-tailed unpaired Student’s t-test.

**Figure S3.**
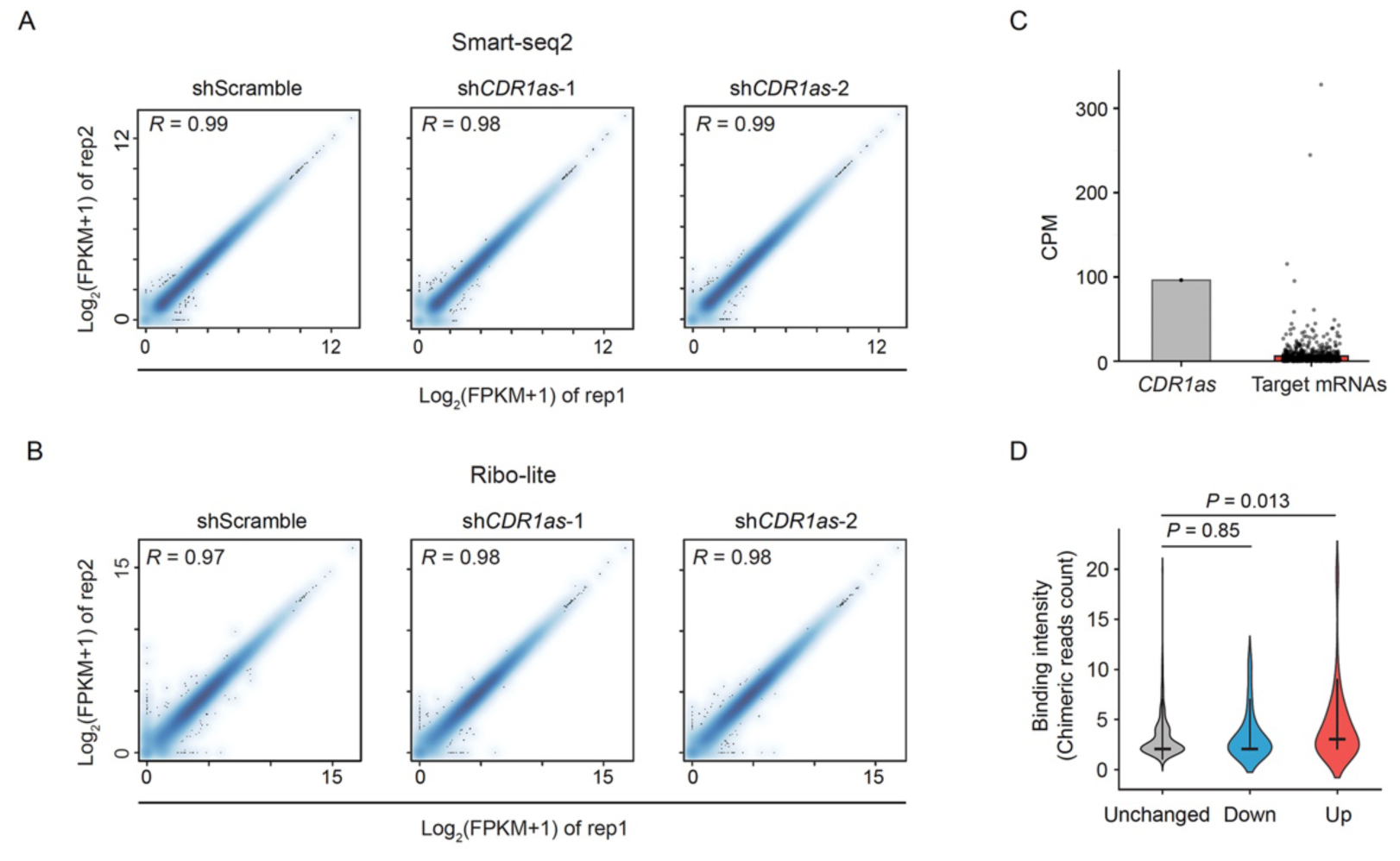
The reproducibility of Smart-seq2 and Ribo-lite datasets. (A-B) Scatter plots showing the reproducibility between two biological replicates in Smart-seq2 (A) and Ribo-lite (B) datasets. (C) Bar plots showing the expression level of *CDR1as* and its target mRNAs. (D) Violin plots showing *CDR1as* binding intensity to target RNAs. The *P*-values are calculated using a Wilcoxon rank-sum test.

**Figure S4.**
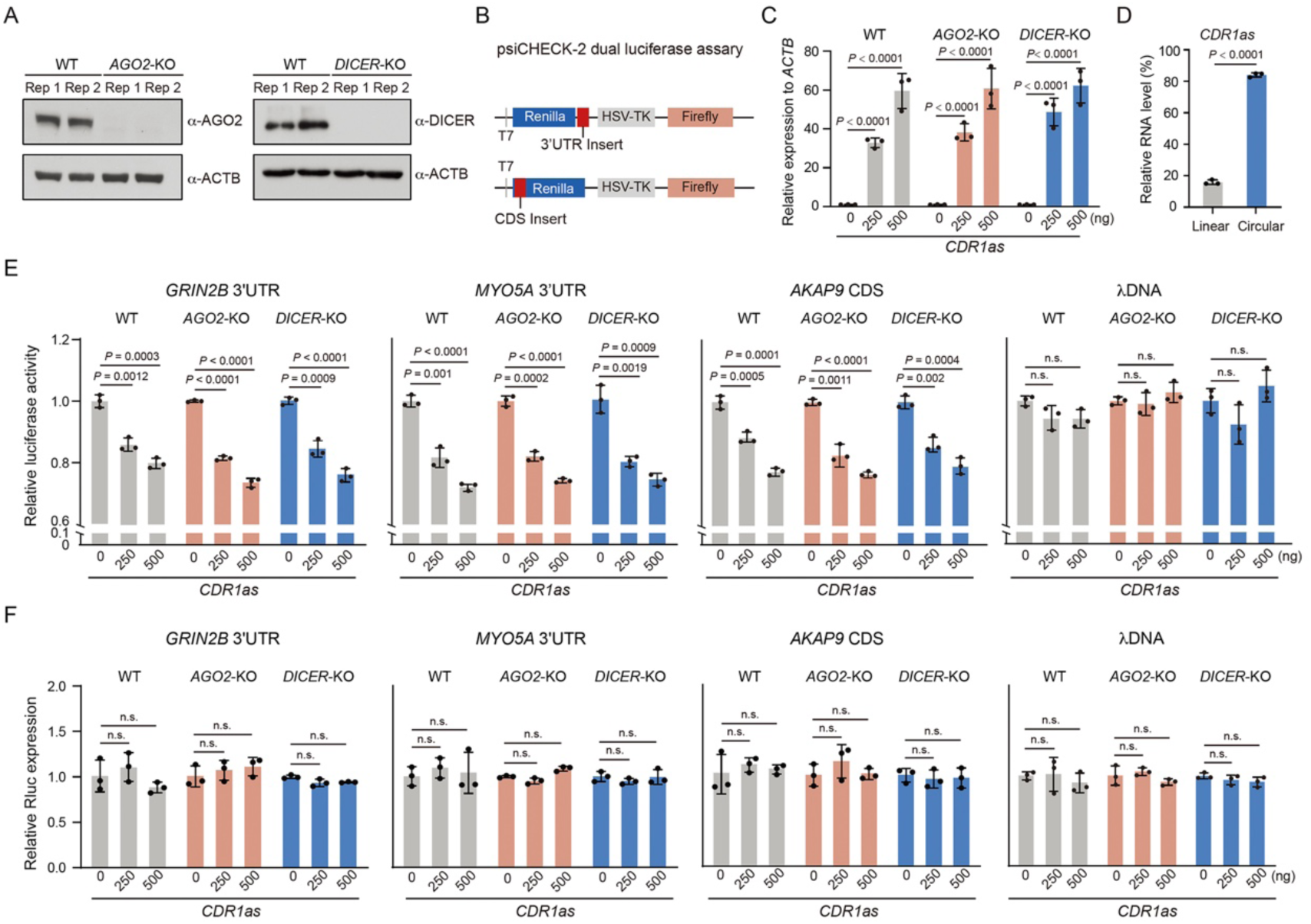
***CDR1as* represses translation independently of AGO2 and miRNAs.** (A) Immunoblotting shows the KO efficiency of AGO2 and DICER in 293T cells. (B) Diagram of dual luciferase reporter assay for characterizing the translational repression of *CDR1as* on target mRNAs. (C) qPCR results showing the expression levels of *CDR1as* in WT, *AGO2*-KO, and *DICER*-KO 293T cells transfected with increasing doses of *CDR1as* plasmid (0, 250, 500 ng). (D) qPCR results showing significantly higher circular-to-linear ratio of *CDR1as* in transfected 293T cells compared to linear transcript. (E) Dual-luciferase assays showing *CDR1as* dose-dependent repression of target mRNA (*GRIN2B*, *MYO5A*, and *AKAP9*) reporters in WT, *AGO2*-KO, and *DICER*-KO 293T cells, with no effect on λDNA control reporter. (F) qPCR results showing unchanged Renilla luciferase reporter mRNA levels after transfection with increasing doses of *CDR1as* plasmid. Data in C, D, E, and F are presented as mean ± s.d.; *n* = 3 independent replicates, two-tailed unpaired Student’s t-test.

**Figure S5.**
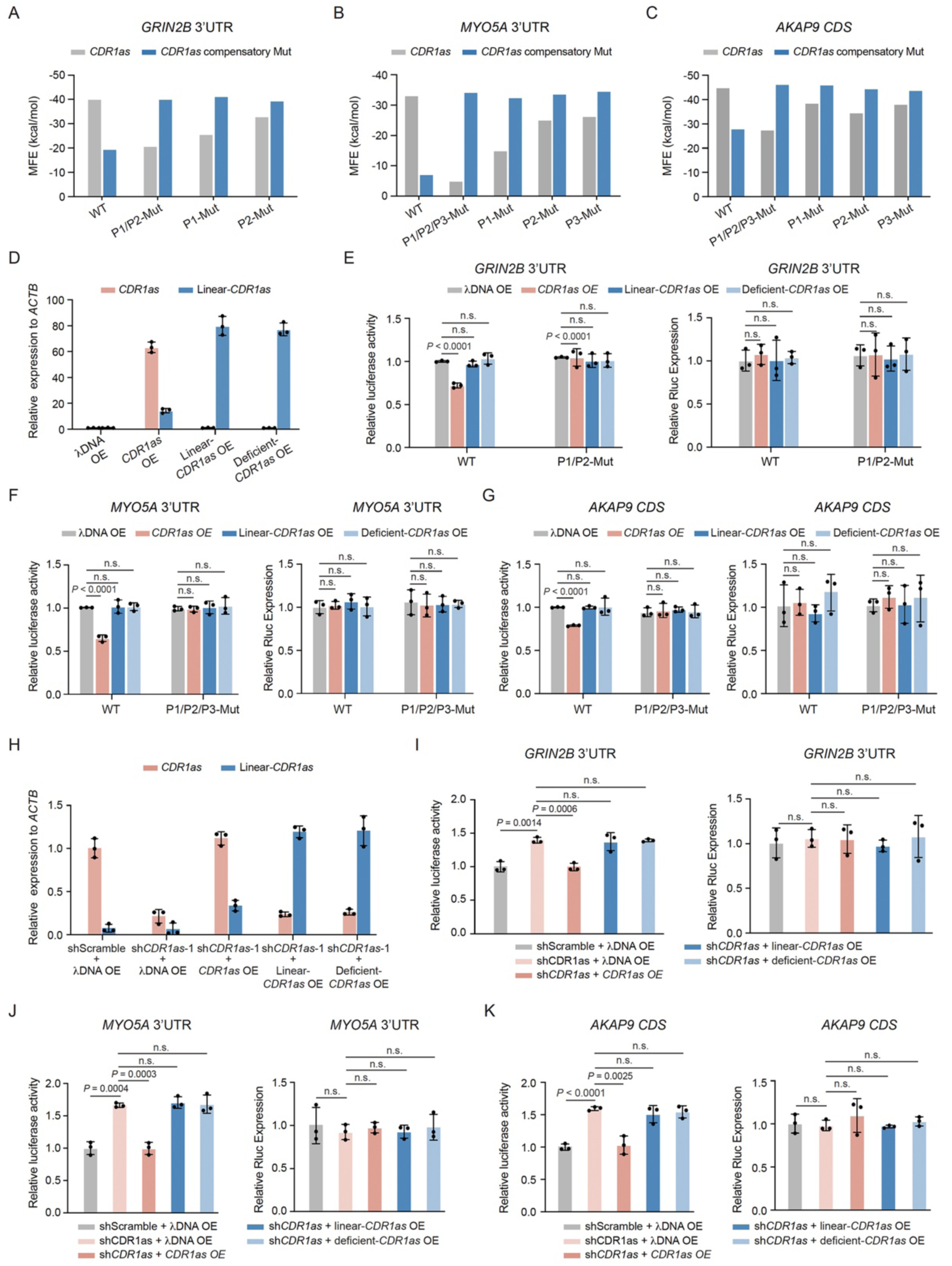
The circular form of *CDR1as* represses target mRNA translation via base-pairing. (A-C) Bar plots showing the MFEs of RNA duplexes formed between *CDR1as* (or its compensatory mutants) and WT or mutated fragments of *GRIN2B* 3’UTR (A), *MYO5A* 3’UTR (B), and *AKAP9* CDS (C). (D) qPCR analysis of *CDR1as* and linear-*CDR1as* expression in circular, linear, or circularization-deficient *CDR1as* (lacking the forward circularization frame) overexpressing HeLa cells. A λDNA vector served as a negative control. (E-G) Luciferase reporter assays showing the relative luciferase activities (left panel) and mRNA levels (right panel) of *GRIN2B* 3’UTR (WT, P1/P2-Mut) reporters (E), *MYO5A* 3’UTR (WT, P1/P2/P3-Mut) reporters (F), *AKAP9* CDS (WT, P1/P2/P3-Mut) reporters (G) in circular, linear, or circularization-deficient *CDR1as* overexpressing HeLa cells. (H) qPCR analysis of *CDR1as* and linear-*CDR1as* expression in *CDR1as* KD and circular, linear, or circularization-deficient *CDR1as* overexpressing 293T cells. (I-K) Luciferase reporter assays showing the relative luciferase activities (left panel) and mRNA levels (right panel) of *GRIN2B* 3’UTR reporter (I), *MYO5A* 3’UTR reporter (J), *AKAP9* CDS reporter (K) in *CDR1as* KD and circular, linear, or circularization-deficient *CDR1as* overexpressing 293T cells. Data in D-K are presented as mean ± s.d.; *n* = 3 independent replicates, two-tailed unpaired Student’s *t*-test.

**Figure S6.**
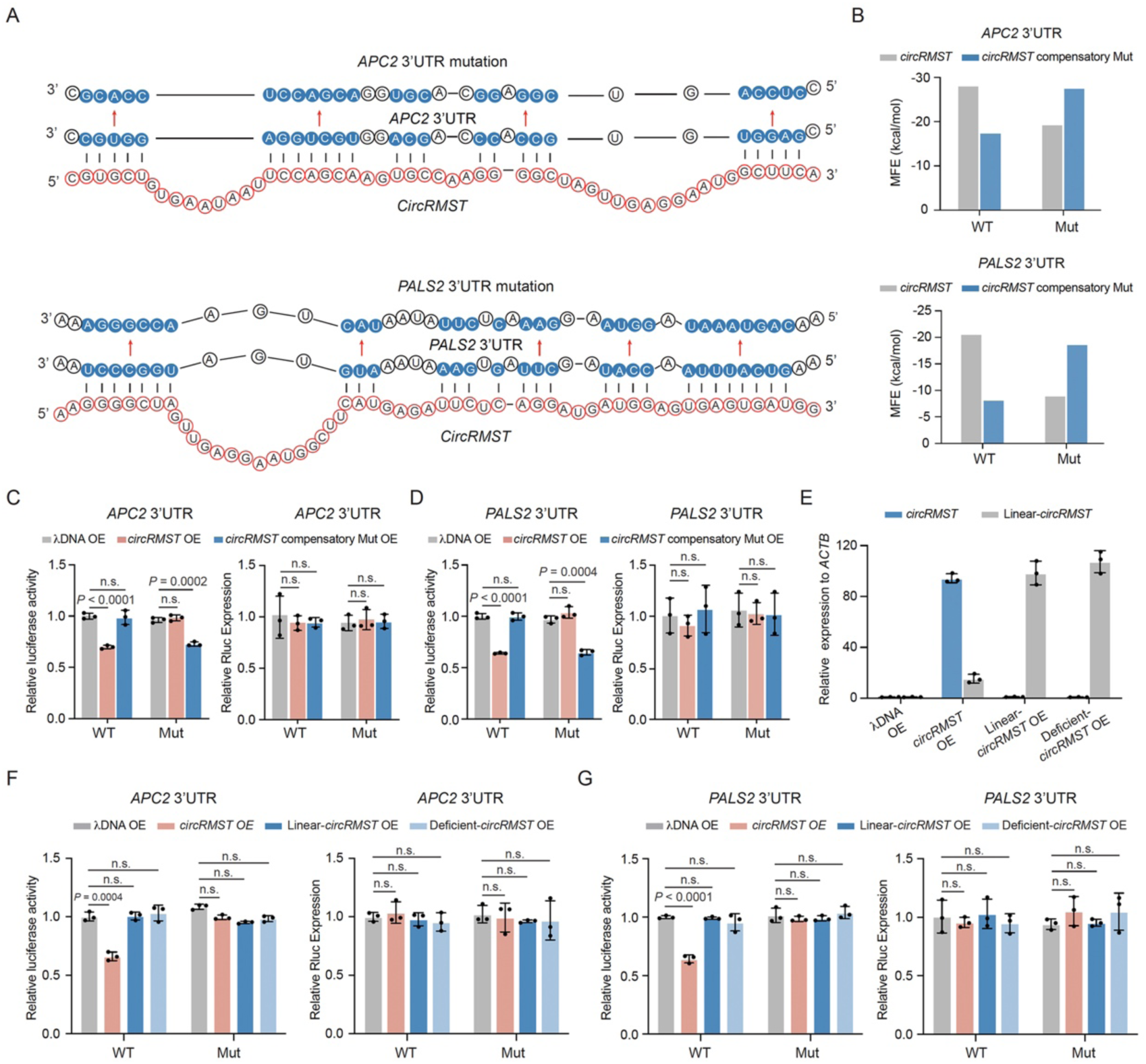
The circular form of *circRMST* represses target mRNA translation via base-pairing. (A) RNA duplexes between *circRMST* (BSJ region) and the interacting fragment of *APC2* or *PALS2* 3’UTR. Arrows indicate the mutation regions of *APC2* and *PALS2* 3’UTRs. (B) Bar plots showing the MFEs of RNA duplexes formed between *circRMST* (or its compensatory mutants) and WT or mutated fragments of *APC2* and *PALS2* 3’UTRs. (C-D) Luciferase reporter assays showing relative luciferase activities (left) and mRNA levels (right) of *APC2* (C) and *PALS2* (D) 3’UTR reporters (WT and Mut) in 293T cells overexpressing *circRMST* or compensatory mutant *circRMST*. (E) qPCR analysis of *circRMST* and linear-*circRMST* expression in 293T cells overexpressing circular, linear, or circularization-deficient *circRMST*. λDNA vector served as a negative control. (F-G) Luciferase reporter assays showing the relative luciferase activities (left panel) and mRNA levels (right panel) of *APC2* 3’UTR (WT and Mut) reporters (F), *PALS2* 3’UTR (WT and Mut) reporters (G) in 293T cells overexpressing circular, linear, or circularization-deficient *circRMST.* Data in C-G are presented as mean ± s.d.; *n* = 3 independent replicates, two-tailed unpaired Student’s *t*-test. n.s., non-significant.

**Figure S7.**
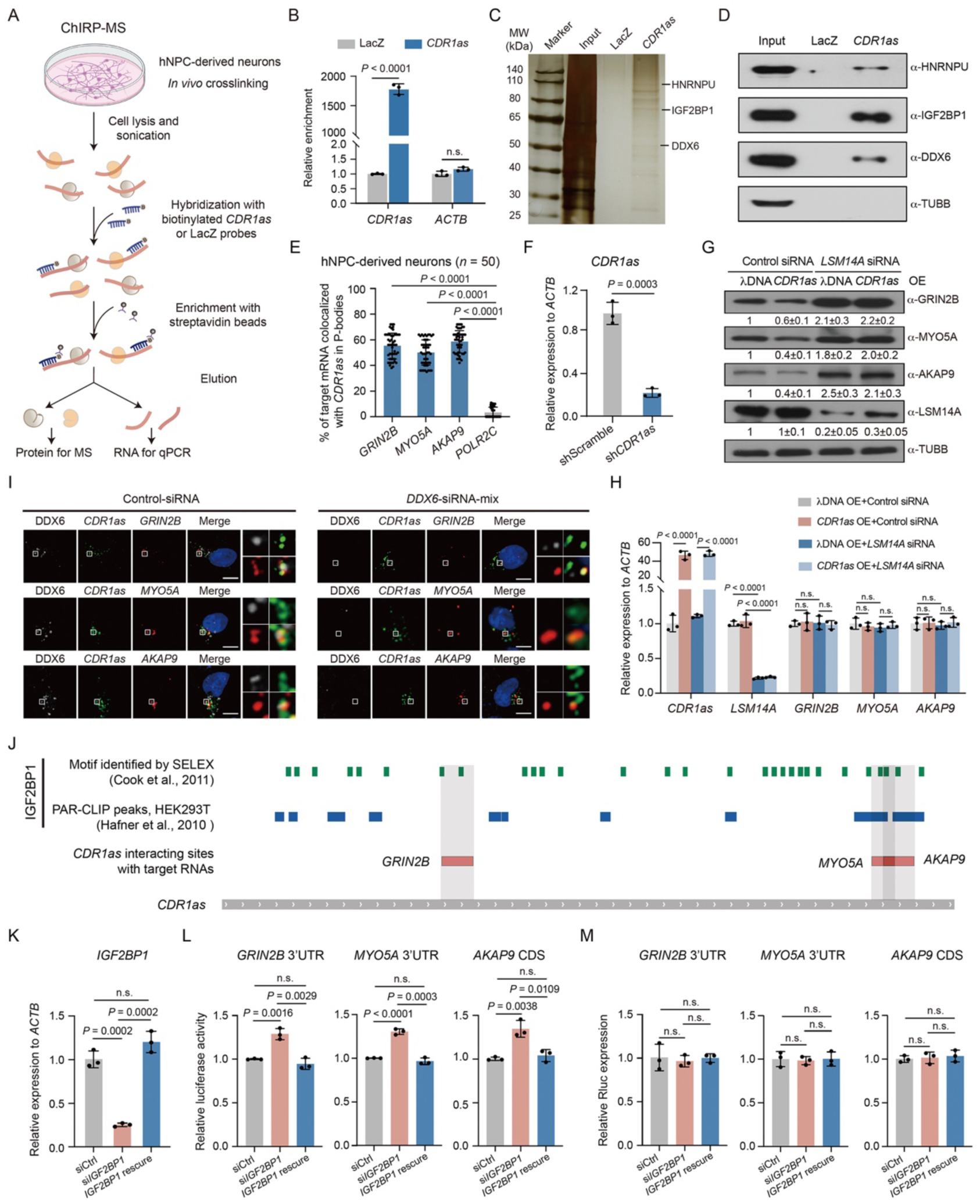
The interactions between *CDR1as* and P-body proteins. (A) Schematic illustrating the ChIRP-MS strategy for detecting *CDR1as*-interacting proteins using *CDR1as* and LacZ control probes in neurons. (B) qPCR results show the enrichment of *CDR1as* in *CDR1as* ChIRP compared to LacZ ChIRP. *ACTB* serves as a negative control. (C) Silver staining shows *CDR1as* ChIRP-enriched RBPs. (D) Immunoblotting shows that the P-body proteins (HNRNPU, IGF2BP1, and DDX6) are enriched in *CDR1as* ChIRP compared to LacZ ChIRP controls. TUBB serves as a negative control. (E) The percentage of colocalization between *CDR1as* and targets (*GRIN2B, MYO5A,* and *AKAP9*) in P-bodies (n = 50 neurons per group). *POLR2C* serves as a non-target control. Data are presented as mean ± s.d.; two-tailed unpaired Student’s *t*-test. (F) qPCR showing *CDR1as* knockdown efficiency using the pTRIPZ shRNA system in neurons. shScramble RNA served as a negative control. (G) Immunoblotting shows *CDR1as* targets and LSM14A protein levels after *LSM14A* knockdown in *CDR1as*-overexpressing neurons. Band intensities were normalized to control siRNA-treated λDNA overexpressed neurons; *n* = 3 independent replicates. (H) qPCR shows *CDR1as*, target mRNAs, and *LSM14A* RNA levels after *LSM14A* knockdown in *CDR1as*-overexpressing neurons. (I) IF/smFISH showing colocalization of *CDR1as* with target mRNAs (*GRIN2B*, *MYO5A*, and *AKAP9*) in *DDX6* knockdown and control neurons. DDX6, gray; DAPI, blue; *CDR1as*, green; target mRNAs, red. Scale bar, 5 μm.(J) Schematic of IGF2BP1 binding sites on *CDR1as*. The illustration integrates the IGF2BP1 binding motif from Cook *et al.*^51^ with PAR-CLIP peak data from Hafner *et al.*^52^ on the *CDR1as* sequence. (K) qPCR analysis showing *IGF2BP1* expression levels in *IGF2BP1*-knockdown and rescue 293T cells compared with control siRNA-treated cells. (L-M) Luciferase reporter assays showing the relative luciferase activities (L) and mRNA levels (M) of *GRIN2B* 3’UTR, *MYO5A* 3’UTR, and *AKAP9* CDS reporters in *IGF2BP1*-knockdown and *IGF2BP1* rescue 293T cells. Data in B, F, H, K, L, and M are presented as mean ± s.d.; *n* = 3 independent replicates, two-tailed unpaired Student’s *t*-test. n.s., non-significant.

**Figure S8.**
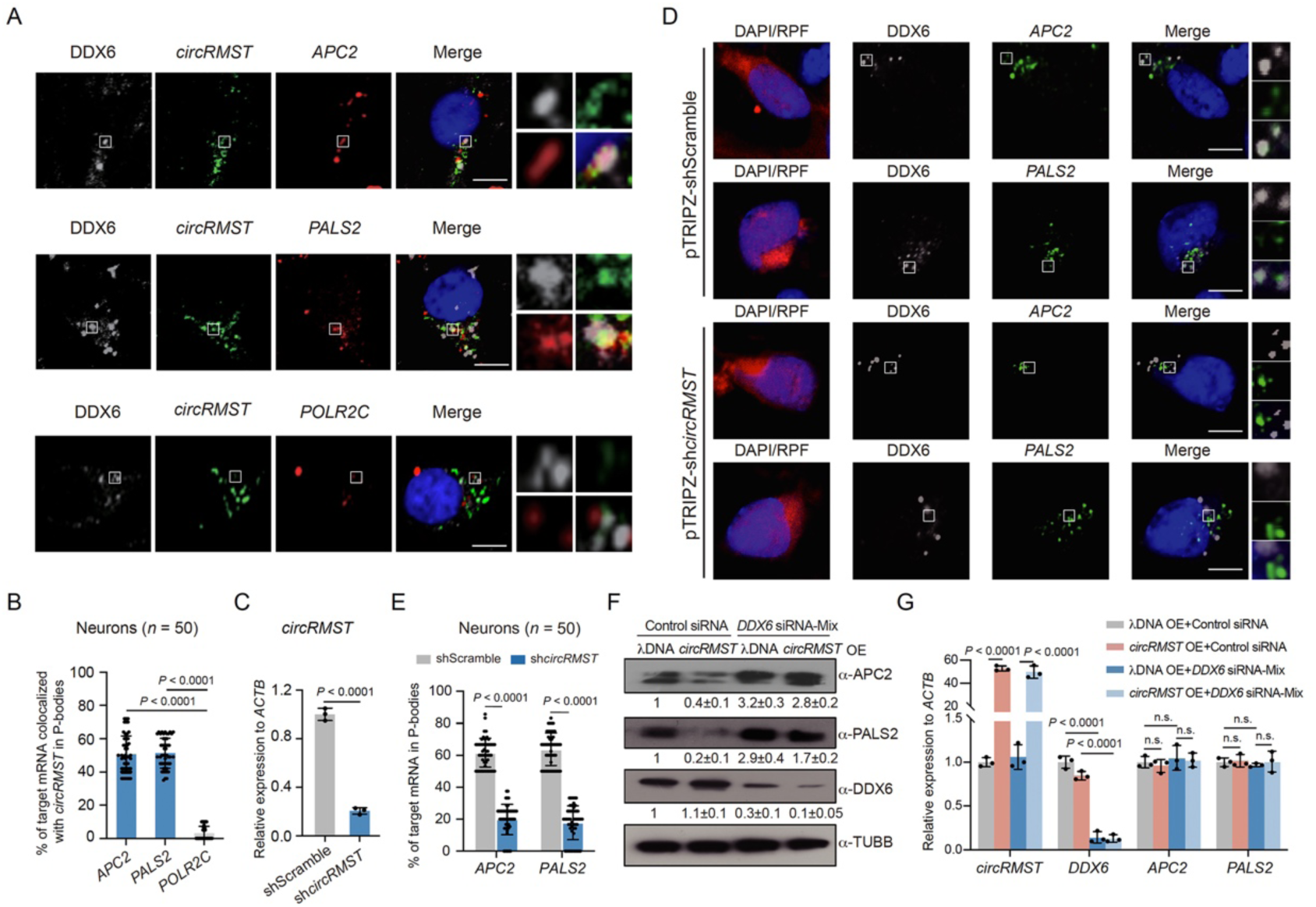
*CircRMST* sequesters target mRNAs into P-bodies for translational repression. (A) IF/HCR-FISH showing colocalization of *circRMST* with its target mRNAs (*APC2* and *PALS2*) and the P-bodies marker protein DDX6 in neurons. *POLR2C* shows no colocalization. DDX6, gray; DAPI, blue; *CDR1as*, green; target mRNAs, red. Scale bar, 5 μm. (B) The percentage of colocalization between *circRMST* and targets (*APC2* and *PALS2*) or non-target *POLR2C* in P-bodies (*n* = 50 neurons per group). Data are presented as mean ± s.d.; two-tailed unpaired Student’s *t*-test. (C) qPCR showing the expression of *circRMST* in the pTRIPZ-induced *circRMST* knockdown neurons. shScramble serves as a negative control. (D) IF/HCR-FISH showing *circRMST* target mRNAs localized in P-bodies within shScramble-treated neurons but not in *circRMST* knockdown neurons (tRFP+, red). Scale bar, 5 μm. (E) Percentage of *circRMST* target mRNAs localized in P-bodies (*n* = 50 neurons) in *circRMST* knockdown versus shScramble neurons. Data are presented as mean ± s.d.; two-tailed unpaired Student’s *t*-test. (F) Immunoblotting shows *circRMST* targets and DDX6 protein levels after *DDX6* knockdown in *circRMST*-overexpressing neurons. Band intensities were normalized to control siRNA-treated λDNA overexpressed neurons; *n* = 3 independent replicates. (G) qPCR shows *circRMST*, target mRNAs, and *DDX6* RNA levels after *DDX6* knockdown in *circRMST*-overexpressing neurons. Data in C and G are presented as mean ± s.d.; *n* = 3 independent replicates, two-tailed unpaired Student’s *t*-test. n.s., non-significant.

**Figure S9.**
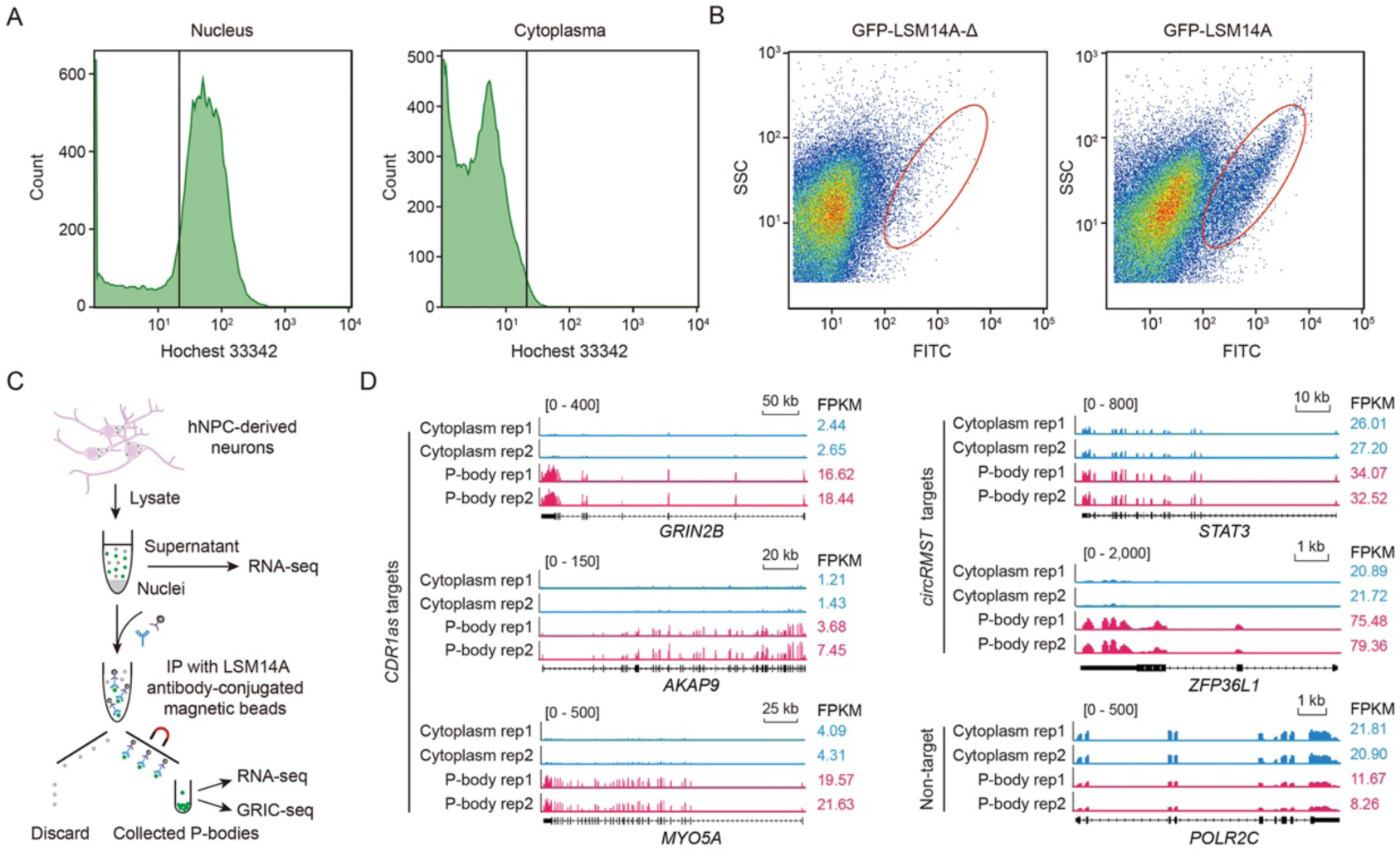
CircRNAs and their target mRNAs are enriched in purified P-bodies from neurons. (A) Flow cytometry plots showing the counts of Hochest33342-stained nuclei and cytoplasmic particles by fluorescence intensity. (B) Flow cytometry plots illustrating P-body purification with cell sorter gating for GFP-LSM14A-positive P-bodies, using GFP-LSM14A-Δ particles as a negative control. (C) Schematic illustrating the experimental procedure for immunoprecipitation of P-bodies using the LSM14A antibody in hNPC-derived neurons. The purified P-bodies were subsequently used for RNA-seq and GRIC-seq analysis. (D) RNA-seq tracks showing the expression of *CDR1as* target mRNAs (*GRIN2B, AKAP9, and MYO5A)*, *circRMST* target mRNAs (*STAT3* and *ZFP36L1*) and the non-target control *POLR2C* in the cytoplasm and P-bodies of hNPC-derived neurons. FPKM values are shown on the right side. Rep, replicates.

